# Repeat element–anchored enhancer–promoter interactions (REEPIs) shape the regulatory landscape of hormone-resistant breast cancer

**DOI:** 10.64898/2026.09.09.750334

**Authors:** Xufeng Shu, Shengliang Ni, Mitsuyoshi Murata, Maierdan Palihati, Hisashi Miura, Ichiro Hiratani, Chi Wai Yip, Hazuki Takahashi, Takeya Kasukawa, Noriko Saitoh, Masaki Kato, Piero Carninci

## Abstract

Repeat elements (REs) comprise over half of the human genome and are increasingly recognized as regulatory components, yet their role in therapy resistance remains unclear. Here, we integrate CAGE and RADICL-seq to map regulatory elements and RNA–chromatin interactions in estrogen receptor– positive breast cancer cells (MCF7) and their long-term estrogen-deprived (LTED) derivatives. We identify repeat element–anchored enhancer–promoter interactions (REEPIs), defined by the presence of annotated REs at both enhancer and promoter sites. In LTED cells, REEPIs are enriched for ERV1 and ERVL elements and involve specific repeat pairings associated with transcription factor combinations such as SP1–SMAD3 and SP1–NFYC. At the gene level, these REEPIs are preferentially linked to genes involved in chromatin remodeling, DNA repair, and lineage plasticity. Our findings reveal a repeat-guided regulatory architecture associated with transcriptional adaptation in endocrine resistance and identify REEPIs as a previously underexplored component of regulatory reprogramming in hormone-resistant breast cancer.

## Introduction

Repeat elements (REs), which account for more than half of the human genome, are primarily derived from transposable element (TE) remnants and short tandem repeats (STRs)^1^. Once considered “junk” DNA, REs are now recognized as dynamic components of gene regulation^2^. Many have retained enhancer- or promoter-like activities and can modulate transcription and chromatin structure in both physiological and pathological contexts^3,4^. TEs are broadly categorized into retrotransposons— including long terminal repeat (LTR) elements, long interspersed nuclear elements (LINEs), and short interspersed nuclear elements (SINEs)—and DNA transposons such as Helitrons and hATs^5^. Although most TEs have lost their ability to mobilize, they frequently function as cis-regulatory elements during development and in disease states^6^. Emerging evidence implicates aberrant TE activation in cancer, where TE-derived promoters and enhancers can initiate oncogene transcription, rewire gene regulatory networks, and reshape chromatin landscapes^7,8^. However, the full extent to which REs contribute to transcriptional regulation in cancer remains insufficiently characterized.

Estrogen receptor (ER)-positive breast cancer, a major clinical subtype, initially depends on estrogen signaling but often acquires resistance to endocrine therapies^9^. MCF7 cells serve as a well-established ER-positive model, and their long-term estrogen-deprived (LTED) derivatives mimic features of hormone-independent growth and therapy resistance^10^. While genetic and epigenetic alterations have been implicated in this transition, the regulatory role of REs under estrogen-deprived conditions remains largely unexplored.

To address this gap, we systematically mapped cis-regulatory elements (CREs) and enhancer– promoter interactions (EPIs) in MCF7 and LTED cells using cap analysis of gene expression (CAGE)^11^ and RNA and DNA interacting complexes ligated and sequenced (RADICL-seq)^12^. CAGE enables high-resolution identification of active promoters and enhancers, whereas RADICL-seq captures genome-wide RNA–chromatin interactions. Integrating CAGE-defined regulatory elements with RADICL-seq contacts enables the inference of candidate enhancer–promoter interactions mediated by promoter- and enhancer-associated RNAs. By integrating these datasets, we constructed a comprehensive regulatory landscape of EPIs for both estrogen-responsive and hormone-deprived states.

Our integrated analysis revealed that REs are extensively involved in EPIs under both estrogen-responsive and hormone-deprived conditions. In particular, we identified a distinct subset of interactions anchored by repeat elements at both ends, suggesting that REs may provide sequence and structural features that facilitate regulatory communication. These interactions were linked to genes involved in chromatin remodeling, DNA repair, and lineage plasticity—processes associated with endocrine resistance. Together, our findings highlight a previously underappreciated role for REs in shaping transcriptional networks in hormone-resistant breast cancer and suggest that repeat element– guided regulatory interactions may contribute to therapeutic escape mechanisms.

## Results

### Comprehensive identification of CREs and EPIs in MCF7 and LTED cells

We generated CAGE libraries from MCF7 and LTED cells and applied the SCAFE pipeline^13^ to systematically define active cis-regulatory elements (CREs), including promoters and enhancers. This analysis identified 32,165 CREs in MCF7 (24,777 promoters; 7,388 enhancers) and 34,020 CREs in LTED cells (25,353 promoters; 8,667 enhancers) (Fig. 1a).

**Fig. 1:**
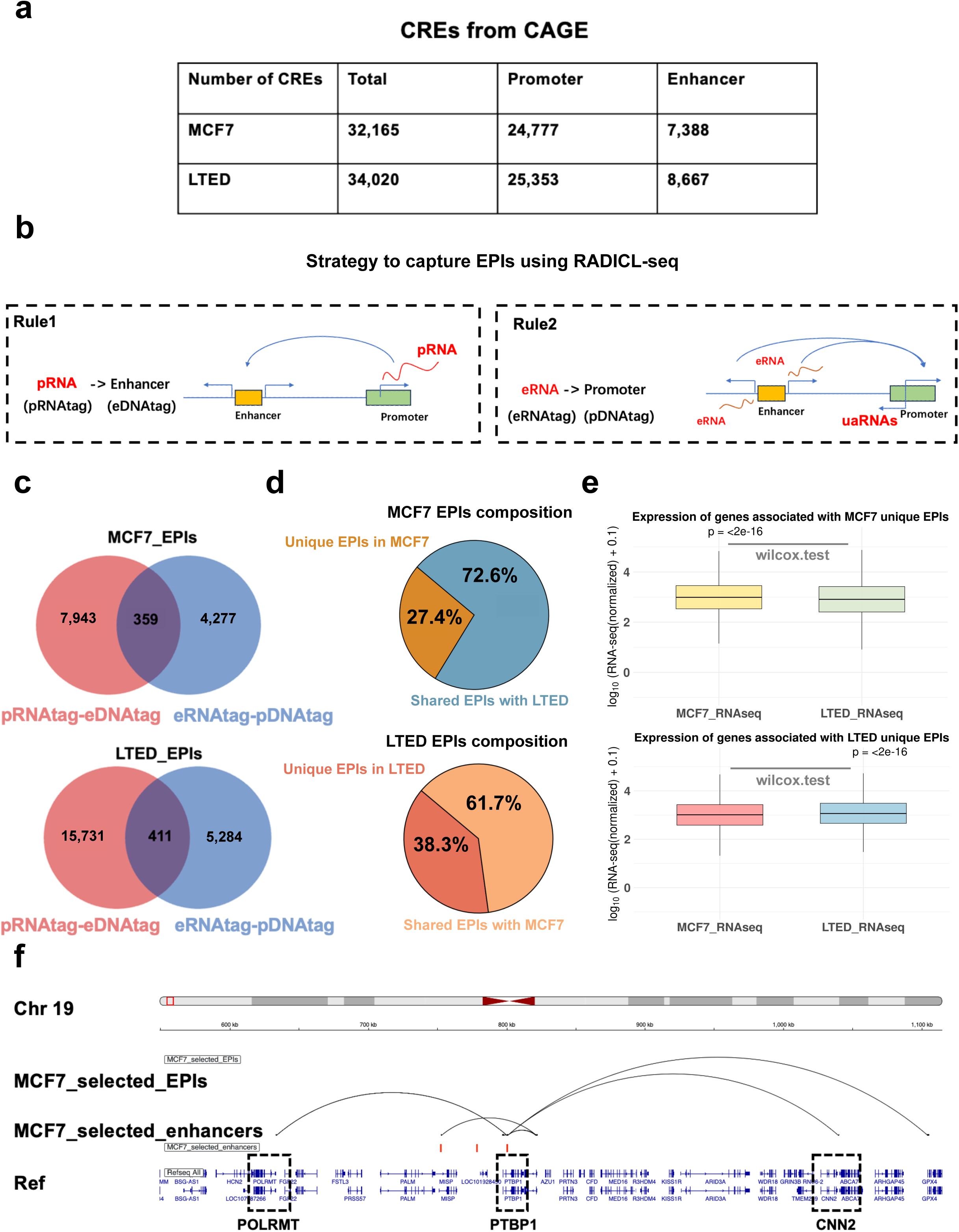
Integrative identification of cis-regulatory elements (CREs) and enhancer–promoter interactions (EPIs) in MCF7 and LTED cells. **a,** CAGE profiling was used to define active CREs in MCF7 and LTED breast cancer cells. Promoters and enhancers were annotated using the SCAFE pipeline. b, Overview of the EPI identification strategy. Filtered RADICL-seq data were integrated with CAGE-defined CREs, and EPIs were categorized into two classes: Rule 1, involving interactions between promoter-derived RNAs (pRNAs) and enhancer DNA (pRNA–eDNA); and Rule 2, involving interactions between enhancer RNAs (eRNAs) and promoter DNA (eRNA–pDNA). c, Overlap between EPIs defined by Rule 1 (pRNA– eDNA) and Rule 2 (eRNA–pDNA) in MCF7 (n = 12,579) and LTED (n = 21,426) cells. d, Proportion of EPIs that are cell type–specific or shared between MCF7 and LTED cells. e, Cell type–specific EPIs are associated with significantly higher expression of their target genes in the corresponding cell type. f, Representative EPIs near the PTBP1 locus, showing concordance with previously reported RNA-associated contacts. The CAGE-defined enhancers (orange bars) engage in EPIs (black arcs) with the promoters of *POLRMT*, *PTBP1*, and *CNN2* (highlighted by dashed boxes).

Next, to map enhancer–promoter interactions (EPIs), we integrated CAGE-defined CRE coordinates with RNA–chromatin contacts detected by RADICL-seq. RADICL-seq captures RNA–DNA chimeras formed by spatial proximity, reflecting RNA contacts associated with both canonical chromatin loops and more transient RNA-mediated interactions. We restricted the analyses to significant RNA–DNA contacts (adjusted P ≤ 0.001), which accounted for 9.85% and 10.99% of chimeric reads in MCF7 and LTED cells, respectively. We operationally defined RNA-supported contacts linking CAGE-defined promoters and enhancers as enhancer–promoter interactions (EPIs). We then classified these EPIs into two categories based on RNA origin (Fig. 1b): Rule 1: Promoter-derived RNAs (pRNA) interacting with enhancer DNA (eDNA), denoted pRNA–eDNA. Rule 2: Enhancer RNAs (eRNAs) interacting with promoter DNA (pDNA), denoted eRNA–pDNA.

We identified 12,579 unique EPIs in MCF7 (8,302 Rule 1; 4,636 Rule 2) and 21,426 EPIs in LTED (16,142 Rule 1; 5,695 Rule 2) (Fig. 1c). Only ∼8% overlap was observed between Rule 1 and Rule 2 EPIs, indicating that Rule 1 and Rule 2 capture distinct modes of RNA-associated enhancer–promoter communication. We next compared the genomic distance distributions of Rule 1 and Rule 2 EPIs. Rule 1 EPIs spanned significantly greater genomic distances than Rule 2 EPIs in both cell types (Supplementary Fig. 1), consistent with broader detection from stable promoter transcription relative to the typically low and short-lived output of eRNAs. Accordingly, the sensitivity of Rule 1 depends on RNA abundance and stability, and Rule 1 is particularly informative for long-range contacts. Another key advantage of RADICL-seq is its strand-independent capture of DNA, which allows identification of RNA–DNA interactions regardless of DNA strand orientation. This feature enables the detection of enhancer–promoter contacts involving upstream antisense RNAs (uaRNAs or PROMPTs) transcribed from bidirectional promoters (Fig. 1b). Many uaRNAs and eRNAs harbor Alu repeat elements, and recent studies have shown that complementary Alu sequences mediate enhancer– promoter specificity via direct RNA–RNA pairing^14^ (Supplementary Fig. 2). Consistently, de novo motifs enriched at EPI-associated promoters and enhancers showed significant clustering within the AluSx consensus sequence in both MCF7 and LTED cells (Supplementary Figs. 3 and 4), supporting a model of sequence-guided, RNA-mediated enhancer–promoter communication.

We also observed a 1.7-fold increase in total regulatory interactions in LTED cells relative to MCF7 (Fig. 1c), with a marked expansion of Rule 1 interactions. This expansion reveals a broader landscape of RNA-associated enhancer–promoter interactions in LTED cells, consistent with extensive transcriptional reprogramming associated with endocrine resistance. To assess the specificity of EPIs, we compared the RADICL-seq–derived EPI sets identified in MCF7 and LTED cells. Despite the distinct hormonal contexts of these models, the majority of EPIs were shared, with only 27.4% and 38.3% being unique to MCF7 and LTED, respectively (Fig. 1d). Given that EPIs are frequently associated with transcriptional activation, we next examined the relationship between cell-specific EPIs and gene expression. Genes linked to MCF7-specific EPIs exhibited significantly higher expression in MCF7 cells than in LTED cells, and vice versa (Fig. 1e), suggesting that condition-specific EPIs are associated with cell state-dependent transcriptional programs.

A notable example was observed at the *PTBP1* locus. In MCF7 cells, an enhancer located within an intronic region of the *PTBP1* gene on chromosome 19 engaged in EPIs with the promoters of *PTBP1, POLRMT* and *CNN2* genes (Fig. 1f). This locus was selected because similar interactions were previously reported by RIC-seq analyses, albeit in different cell types^14^, supporting the biological relevance and reproducibility of these regulatory contacts.

Together, these findings demonstrate that integrating CAGE with RADICL-seq enables systematic identification of RNA-supported enhancer–promoter interactions and their association with cell state– specific transcriptional programs in hormone-sensitive and endocrine-resistant breast cancer cells.

### Characteristics of RADICL-seq–derived EPIs

We next characterized the genomic architecture of EPIs identified by RADICL-seq. Notably, a substantial fraction of interactions occurred in *trans*—between different chromosomes—accounting for ∼54% in MCF7 and ∼60% in LTED cells (Fig. 2a). Given the transient nature and limited reproducibility of *trans* interactions reported for proximity ligation-based assays^15–17^, subsequent analyses were restricted to *cis* interactions occurring within the same chromosome, which are generally more reproducible. Among *cis*-EPIs, the majority spanned less than 1 Mb (∼82.6% in MCF7; ∼85.3% in LTED), consistent with the typical size of topologically associating domains (TADs) in the human genome^18^. Despite this spatial constraint, only ∼32% of enhancers interacted with their nearest promoter (Fig. 2b), indicating that linear genomic proximity is not a reliable predictor of enhancer– promoter communication. All subsequent analyses focused on these *cis*-EPIs, hereafter referred to simply as EPIs.

**Fig. 2:**
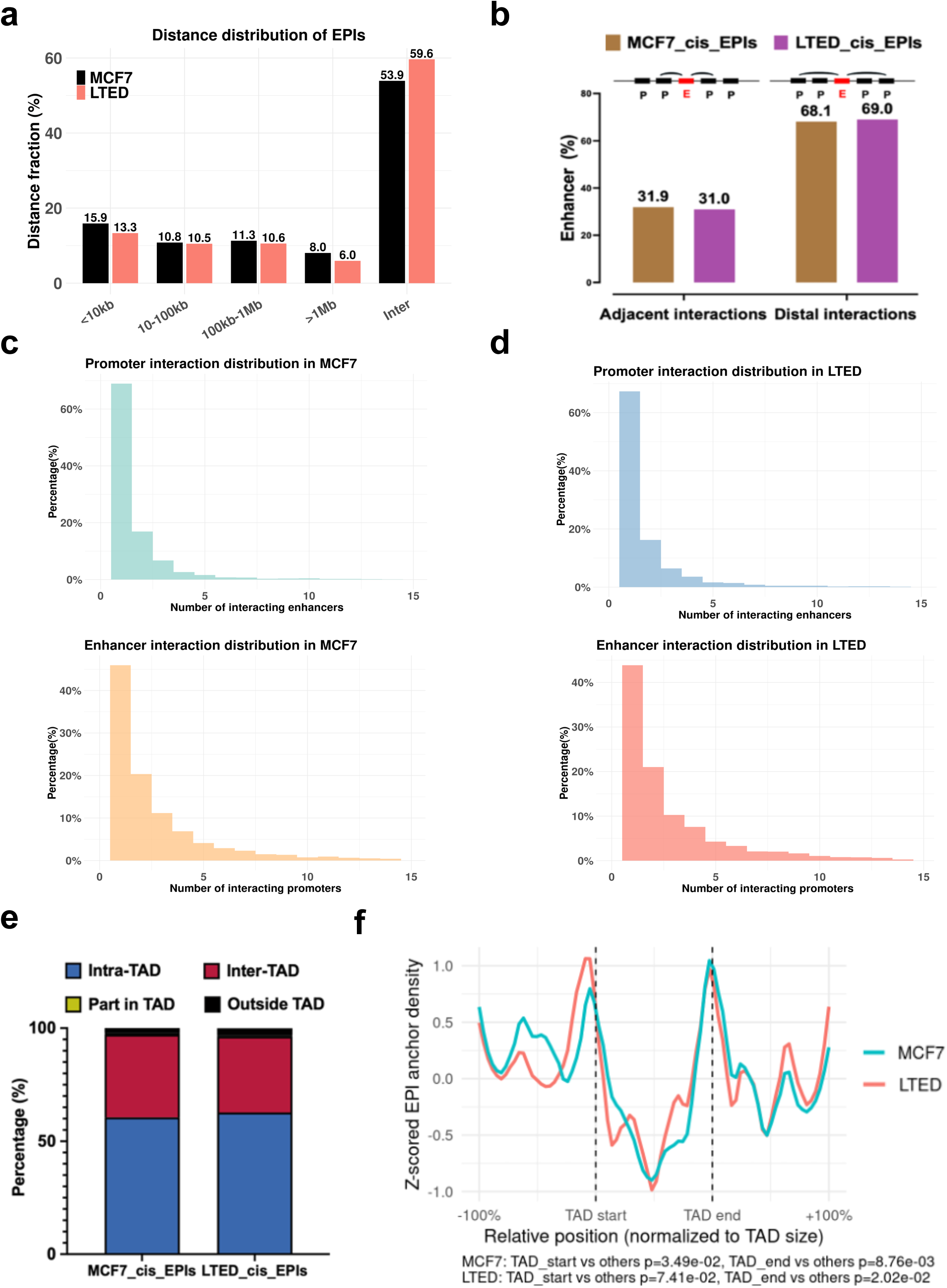
Genomic and topological characteristics of EPIs. **a**, Classification of EPIs based on genomic distance. A substantial fraction of EPIs were interchromosomal (*trans*) in both MCF7 (∼54%) and LTED (∼60%) cells. **b**, Proportion of *cis*-EPIs in which enhancers interact with their nearest annotated promoter. **c**, Top: Distribution of the number of enhancers interacting with each promoter in MCF7 cells. Bottom: Distribution of the number of promoters contacted by each enhancer. **d**, Equivalent analysis in LTED cells. Top: Distribution of enhancer contacts per promoter. Bottom: Distribution of promoter contacts per enhancer. **e,** Classification of EPIs with respect to TAD structures identified from Hi-C data at 25 kb resolution. EPIs were classified into four categories: “Intra-TAD”(blue), in which both anchors were located within the same TAD; “Inter-TAD”(red), in which the two anchors resided in different TADs; “Part in TAD”(yellow), in which only one anchor was located within an annotated TAD while the other lay outside any annotated TAD; and “Outside TAD”(black), in which neither anchor was located within an annotated TAD. **f**, Meta-profile of EPI anchor positions relative to TAD architecture. Each TAD was symmetrically extended by one TAD length upstream and downstream and divided into 300 equal bins (100 upstream, 100 within, 100 downstream). The midpoint of each EPI anchor was mapped to the corresponding normalized bin. Density profiles were Z-score normalized and smoothed using a 3-bin rolling average. Boundary enrichment was statistically evaluated by comparing average Z-scores in boundary bins (±10% from TAD start/end) to “Other” bins using a two-tailed Student’s t-test.

RADICL-seq further revealed a many-to-many interaction landscape, with individual promoters contacted by multiple enhancers and vice versa (Fig. 2c,d), supporting a combinatorial model of transcriptional regulation. To investigate the relationship between EPIs and 3D genome architecture, we intersected EPI anchors with TAD annotations derived from Hi-C (Kato et al., manuscript in preparation). The majority of EPIs were confined within TADs (61% in MCF7; 63% in LTED, Fig. 2e, blue), although a substantial fraction crossed TAD boundaries (37% in MCF7; 34% in LTED, Fig. 2e, red), and a small subset (∼2–3%) occurred entirely outside defined TADs (Fig. 2e, black). To test whether EPIs preferentially localize near TAD boundaries, we performed a genome-wide meta-profile analysis by aligning EPI anchor midpoints to normalized TAD-centered windows. This revealed a pronounced enrichment of EPI anchors near both boundaries of TADs in MCF7 and LTED cells (Fig. 2f), suggesting that TAD boundaries may represent preferential sites for EPI anchors.

Together, these findings highlight the influence of 3D chromatin topology in shaping enhancer– promoter communication and organizing the transcriptional regulatory landscape. We next asked whether specific sequence features of EPI anchors, particularly repeat elements, contribute to this regulatory architecture.

### Repeat elements are enriched at enhancer–promoter interaction sites

To examine whether repeat elements contribute to enhancer–promoter specificity, we annotated all EPI anchors using RepeatMasker (http://www.repeatmasker.org). Here, repeat elements (REs) refer to all RepeatMasker-annotated repetitive sequences, including transposable elements (TEs), simple repeats, low-complexity regions, and other annotated repeat classes. Based on repeat element presence at interaction anchors, EPIs were grouped into three categories: (1) non-repeat EPIs, lacking repeats at both anchors; (2) single-end repeat EPIs, containing repeats at either the enhancer or promoter (further classified as promoter-only or enhancer-only); and (3) repeat element-anchored EPIs (REEPIs), with annotated repeats at both anchors (Fig. 3a).

**Fig. 3:**
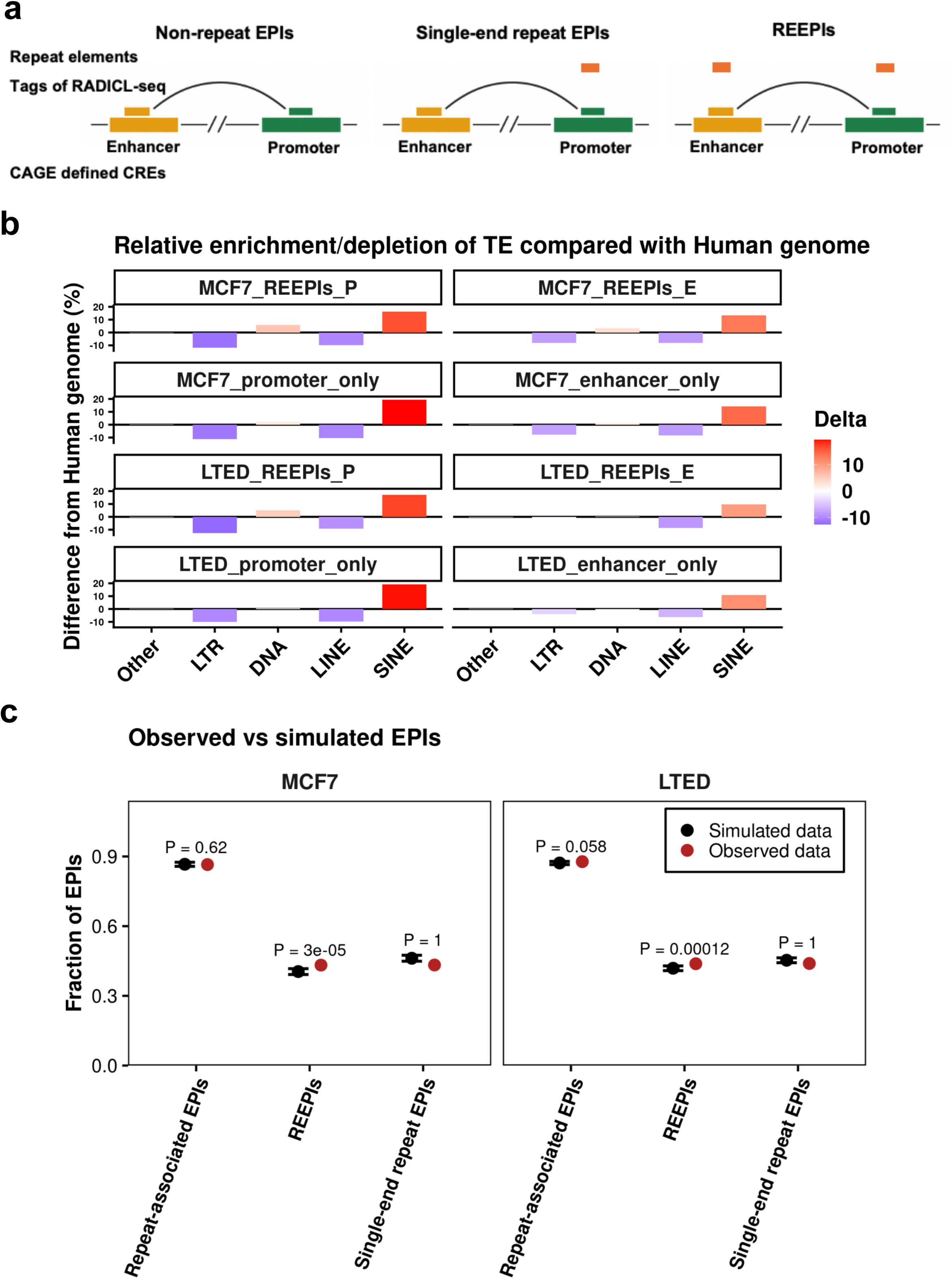
EPIs are enriched for repeat elements. **a**, Schematic classification of RADICL-seq–derived EPIs based on the presence of repeat elements at interaction anchors: (left) non-repeat EPIs, with no repeat elements at either anchor; (middle) single-end repeat EPIs, in which only the enhancer or promoter anchor overlaps a repeat; and (right) repeat element–anchored EPIs (REEPIs), where both anchors overlap annotated repeat elements. **b**, Composition of transposable elements (TEs) at EPI anchors in MCF7 and LTED versus the human genome. Most EPIs contain repeats at one or both anchors (86.5% in MCF7; 87.8% in LTED), with REEPIs forming the largest subset of repeat-associated EPIs (49.97% in MCF7; 49.94% in LTED). TE compositions differ from genome background across categories. Relative enrichment or depletion of TE classes at promoters (P) and enhancers (E) for each EPI class compared with genomic prevalence. Positive values denote over-representation relative to genomic prevalence, with SINEs showing particularly strong enrichment at EPI anchors. **c**, Fractions of repeat-associated EPIs (repeats at one or both anchors), REEPIs, and single-end repeat EPIs are shown in MCF7 (left) and LTED (right). Red points indicate the observed fractions. Black points with error bars represent the simulated background (mean ± 95% CI) from 100,000 chromosome-wise shuffles that preserved each enhancer’s and promoter’s repeat status and the total number of EPIs while randomly shuffling enhancer–promoter pairs. *P* values compare the observed fraction with this background. The overall fraction of repeat-associated EPIs matched the simulated expectation (MCF7: *P* = 0.62; LTED: *P* = 0.058), whereas REEPIs were enriched and single-end repeat EPIs were depleted, indicating a bias toward repeat– repeat pairing beyond what is explained by anchor repeat content alone.

In MCF7 cells, we identified 2,506 REEPIs, 1,463 promoter-only, 1,046 enhancer-only, and 782 non-repeat EPIs. LTED cells showed increased numbers: 3,790 REEPIs, 2,081 promoter-only, 1,718 enhancer-only, and 1,058 non-repeat EPIs. Overall, repeats were present at one or both anchors in 86.5% of MCF7 and 87.8% of LTED EPIs, with REEPIs accounting for approximately half of repeat-associated EPIs (49.97% in MCF7; 49.94% in LTED). These observations suggest that repeat elements are widespread at enhancer–promoter sites and represent a prominent feature of enhancer–promoter interaction networks.

We next analyzed the composition of TE families within repeat-containing EPIs, focusing on REEPIs and comparing them to the genome-wide TE landscape estimated from RepeatMasker annotations. Across all EPI categories, TE compositions differed markedly from the genomic background. SINEs were markedly enriched, whereas LINE and LTR elements were underrepresented. Although SINE enrichment is consistent with their known association with active chromatin, these results indicate that the repeat composition of enhancer–promoter interaction sites differs substantially from overall genomic repeat prevalence. DNA transposons showed a modest enrichment relative to their genomic prevalence (Fig. 3b). Notably, the overrepresentation of SINEs at EPI anchors is consistent with a potential role for Alu elements in sequence- or structure-specific pairing between regulatory regions, as suggested by previous evidence implicating Alu-mediated RNA–RNA pairing in enhancer–promoter communication^14^.

To determine whether high repeat prevalence at enhancer–promoter interactions could be entirely explained by repeat content at individual anchors, we performed a Monte Carlo test comparing the observed EPIs with a chromosome-wise simulated background. This background preserved each enhancer’s and promoter’s repeat status and the total number of EPIs, while randomly shuffling enhancer–promoter pairings. We evaluated three groups: repeat-associated EPIs (repeats present at one or both anchors), REEPIs, and single-end repeat EPIs. *P* values were derived from 100,000 constrained shuffles. The overall fraction of repeat-associated EPIs matched the simulated background (MCF7: *P* = 0.62; LTED: *P* = 0.058), indicating that global repeat prevalence at enhancer–promoter contacts can be explained by anchor repeat content alone. By contrast, REEPIs occurred more frequently than expected, whereas single-end repeat EPIs occurred less frequently. These deviations reveal a bias toward enhancer–promoter contacts with repeats at both anchors. Thus, within enhancer– promoter interaction networks, repeat–repeat pairing is favored beyond background expectation, identifying REEPIs as a preferential configuration of enhancer–promoter contacts (Fig. 3c).

### Repeat subclasses differentially shape REEPIs in endocrine-resistant breast cancer

Although LTR elements were globally underrepresented relative to the genomic background (Fig. 3b), comparison between MCF7 and LTED REEPIs revealed selective enrichment of specific LTR subclasses at LTED enhancer anchors. To delineate how specific repeat subclasses are associated with enhancer–promoter communication in endocrine-resistant breast cancer, we profiled the repeat composition of all REEPIs in MCF7 and LTED cells. ERV1 and ERVL elements were markedly enriched at enhancers of LTED REEPIs, indicating selective remodeling of repeat composition at enhancer anchors during the development of endocrine resistance (Fig. 4a,b).

**Fig. 4:**
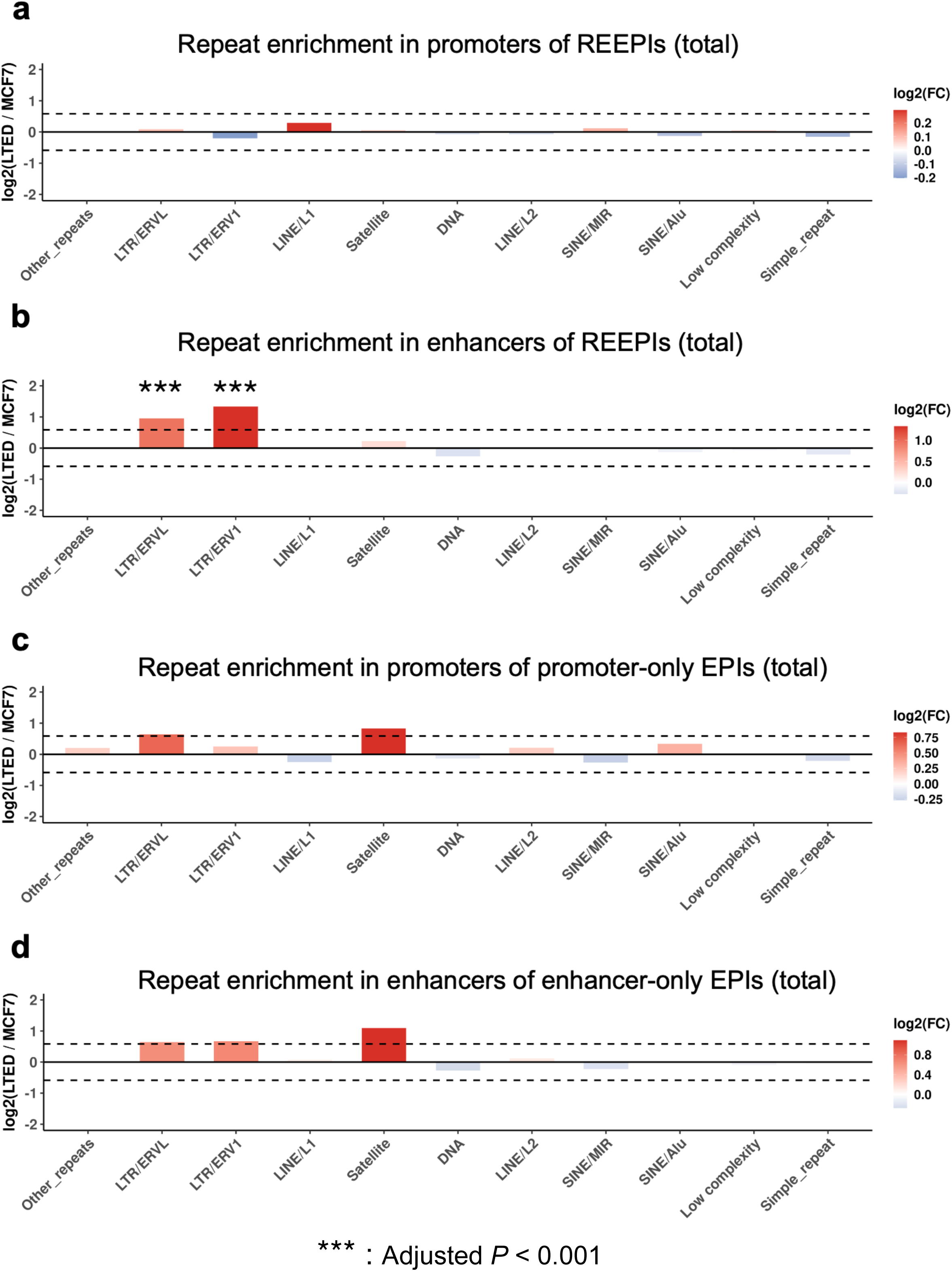
Differential contribution of repeat subclasses to enhancer–promoter interactions in LTED cells. **a**, Enrichment analysis of repeat subclasses at promoter anchors of REEPIs in LTED versus MCF7 cells. **b**, Enrichment of repeat subclasses at enhancer anchors of REEPIs in LTED cells versus MCF7 cells. **c**, Repeat enrichment analysis of promoter-only EPIs. **d**, Repeat enrichment analysis of enhancer-only EPIs.

To determine whether distinct repeat subclasses are preferentially involved across different genomic spans, REEPIs in MCF7 and LTED cells were stratified by linear distance between anchors into <10 kb, 10–100 kb, 100 kb–1 Mb, and >1 Mb (Supplementary Fig. S5). For clarity, interactions shorter than 100 kb were defined as short-range and those longer than 100 kb as long-range. ERVL elements displayed a pronounced long-range preference in LTED, whereas ERV1 elements showed no overall preference when analyzed across all distance classes (two-sided Fisher’s exact test) (Supplementary Fig. S6).

Distance-resolved enrichment analyses confirmed these trends. ERV1 elements in enhancers of LTED REEPIs were significantly enriched at 10–100 kb and >1 Mb, resulting in an overall balanced distribution across distance classes, whereas ERVL elements were preferentially enriched at 100 kb– 1 Mb, consistent with their dominant contribution to long-range interactions (Supplementary Fig. S7). Thus, repeat families exhibit interaction distance–specific preferences, and LTED undergoes a selective remodeling of repeat-associated regulatory contacts. ERV1 and ERVL are broadly enriched across the enhancers of LTED REEPIs, indicating that repeat subclass identity tracks the spatial organization of enhancer–promoter communication. This pervasive enrichment highlights a potential role for these elements in supporting transcriptionally active, eRNA-producing enhancers that participate in regulatory contacts under estrogen-deprived conditions.

We next analyzed single-end repeat EPIs, in which only one anchor harbors a repeat sequence (Fig. 4c,d). In contrast to REEPIs, single-end repeat interactions showed no significant difference between MCF7 and LTED, suggesting that REEPIs—rather than single-end repeat EPIs—more closely associate with the rewired connectivity characteristic of endocrine adaptation.

Together, these findings reveal a selective, distance-dependent reprogramming of repeat element usage within REEPIs in endocrine-resistant breast cancer. The significant enrichment of ERV1 and ERVL elements across a range of interaction distances, coupled with the absence of consistent changes in single-end repeat EPIs, supports the concept that REEPIs represent a prominent repeat architecture associated with the adaptive, transcriptionally permissive enhancer networks that emerge in LTED cells.

### REEPIs define distinct functional programs in endocrine-resistant cells

To evaluate the functional relevance of repeat-associated enhancer–promoter interactions, we performed Gene Ontology (GO) enrichment analysis of genes linked to REEPIs, single-end repeat EPIs (promoter-only and enhancer-only), and non-repeat EPIs in both MCF7 and LTED cells.

In MCF7, single-end repeat and non-repeat EPIs were mainly associated with cell–cell adhesion, morphogenesis, and metabolic processes, consistent with basal epithelial maintenance programs (Supplementary Fig. S8a,c). By contrast, MCF7 REEPIs were significantly enriched for developmental and hormone-responsive pathways, including mammary gland development, epithelial cell differentiation, neurogenesis, and gland morphogenesis (Fig. 5a), reflecting a transcriptional network characteristic of estrogen-dependent epithelial identity. In LTED, single-end repeat and non-repeat EPIs also showed enrichment for RNA metabolic processes (Supplementary Fig. S8b,d), whereas LTED REEPIs were strongly enriched for chromatin- and genome-regulatory functions, including chromatin remodeling, histone modification, regulation of DNA repair, cellular response to hypoxia, and stem cell population maintenance (Fig. 5b). The emergence of these categories suggests that REEPIs are selectively coupled to programs of genome maintenance and adaptive transcriptional plasticity in endocrine-resistant cells.

**Fig 5:**
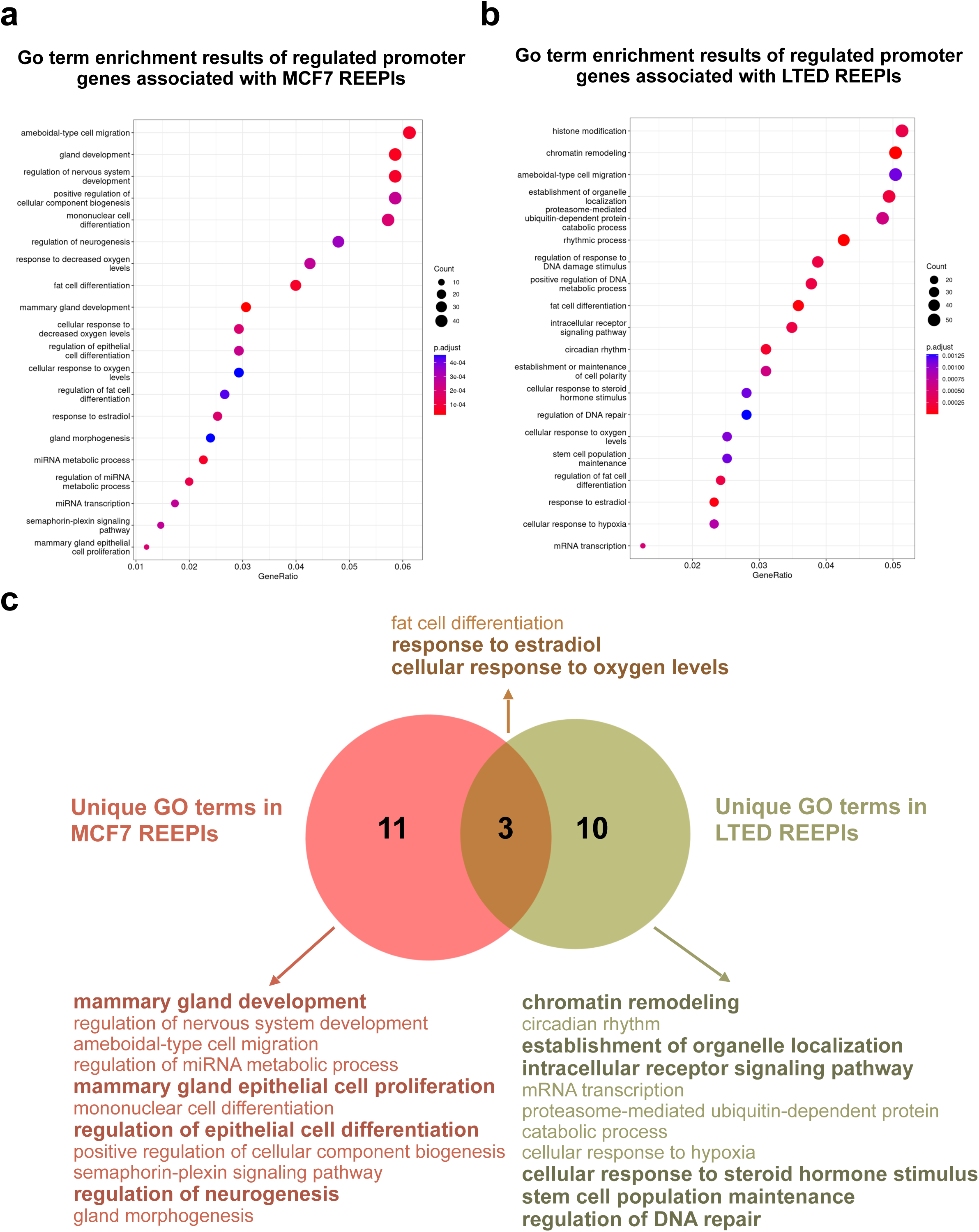
REEPIs are associated with distinct transcriptional reprogramming in LTED cells. **a**, Gene Ontology (GO) enrichment analysis of genes associated with REEPIs in MCF7 cells. **b**, GO enrichment analysis of genes associated with REEPIs in LTED cells. **c**, Venn diagram comparing MCF7 (left) and LTED (right) REEPI-unique GO terms after excluding terms shared with single-end or non-repeat EPIs.

Across both cell states, these comparisons demonstrate that REEPIs display greater functional specificity and coherence than single-end or non-repeat EPIs. To identify the most distinctive programs, we removed GO terms shared with single-end and non-repeat EPIs and then compared the unique terms between MCF7 and LTED REEPIs (Fig. 5c). MCF7-specific REEPI terms were concentrated in mammary gland development, epithelial cell differentiation, and neurogenesis, whereas LTED-specific REEPI terms highlighted chromatin remodeling, regulation of DNA repair, stem cell maintenance, and responses to steroid hormone and oxygen levels. This analysis underscores that, under estrogen-deprived conditions, LTED REEPIs uniquely connect repeat-anchored enhancers to chromatin-restructuring and stress-adaptation pathways characteristic of endocrine resistance.

Collectively, these findings indicate that REEPIs represent a highly specialized and diversified class of enhancer–promoter interactions, linking repeat-encoded regulatory elements to genes governing chromatin dynamics, genome integrity, and hormonal adaptation. This highlights repeat-mediated enhancer architecture as a prominent feature of the transcriptional reprogramming that accompanies the transition from estrogen dependence to endocrine resistance.

### Transcription factor–associated coordination of repeat element pairs may facilitate enhancer– promoter communication in LTED cells

To investigate whether specific “promoter–repeat : enhancer–repeat” pairings contribute to enhancer– promoter interactions in LTED cells, we focused our analysis on REEPIs located outside copy number variation (CNV) regions, as repeat elements have previously been associated with CNV formation and genomic instability^19–21^. Approximately 86% of REEPIs resided outside CNV regions in both MCF7 and LTED cells (MCF7: 2,130 non-CNV vs. 376 CNV; LTED: 3,255 non-CNV vs. 535 CNV; Supplementary Fig. S9a), indicating that the majority of REEPIs are unlikely to be explained by structural genomic alterations.

We next categorized all promoter–enhancer repeat pairings within these non-CNV REEPIs and performed differential interaction analysis between MCF7 and LTED cells. Two pairings were significantly enriched in LTED (*adjusted P* < 0.05, |log₂FC| ≥ 1), including Simple_repeat : LTR/ERV1 and Low_complexity : LTR/ERVL (Fig. 6a). These combinations mirror the previously observed enrichment of ERV1 and ERVL elements in enhancers of LTED REEPIs (Fig. 4b), reinforcing their potential involvement in transcriptional reprogramming under estrogen-deprived conditions. Importantly, the same pairings remained significant even when CNV-overlapping REEPIs were included, indicating that their enrichments were not simply explained by local copy-number amplification (Supplementary Fig. S9b). Because these enriched pairings involve non-homologous repeat families, their interaction is unlikely to depend solely on sequence complementarity or RNA– RNA pairing. As repeat-associated motifs provide candidate binding sites for sequence-specific DNA-binding proteins, we next asked whether transcription factors (TFs) recognizing motifs embedded within distinct repeat types may facilitate these interactions.

**Fig. 6:**
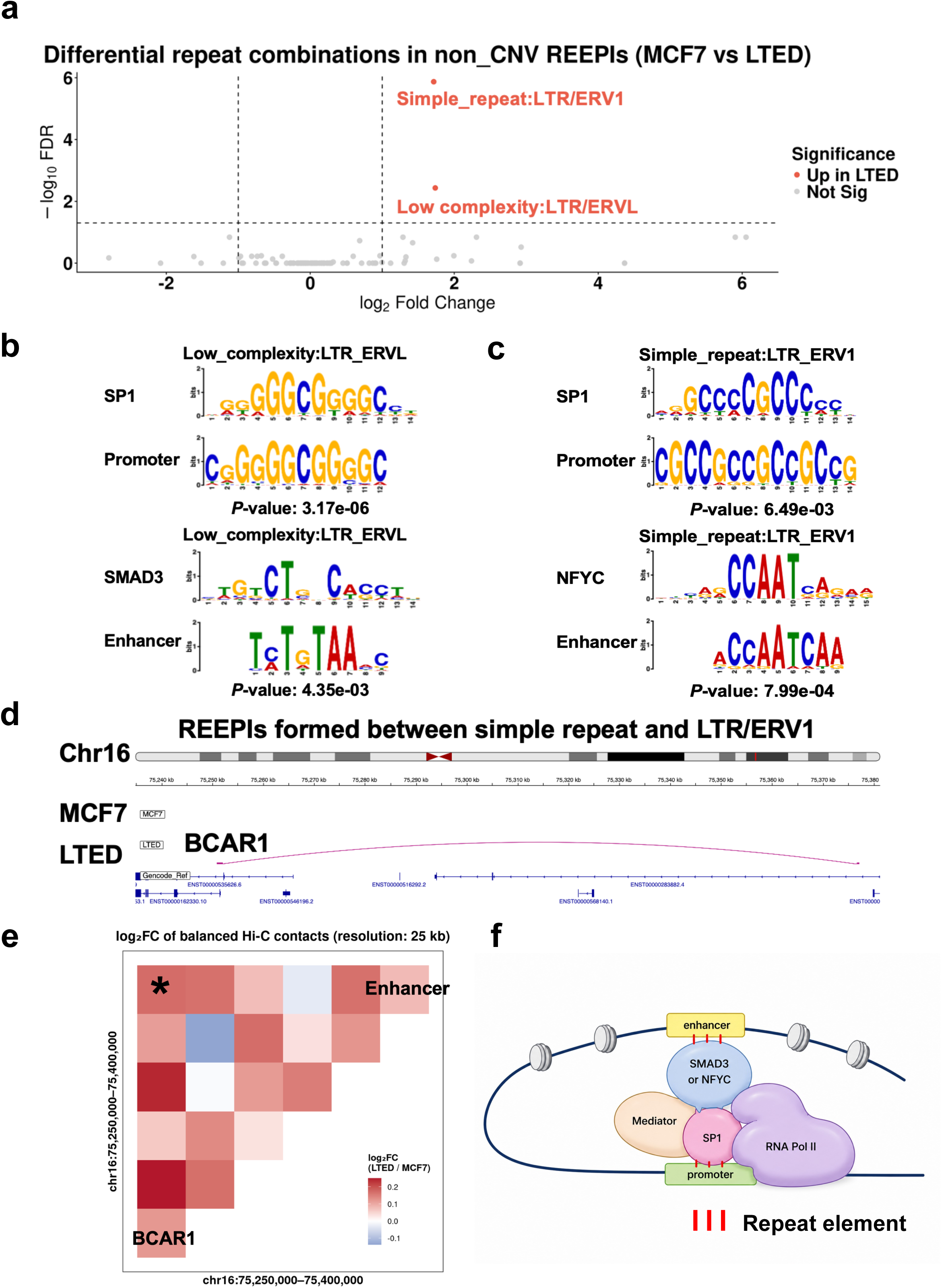
Transcription factor–associated coordination of repeat element pairs may facilitate enhancer–promoter communication in LTED cells. **a**, Volcano plot of promoter–repeat : enhancer–repeat pairings (LTED vs MCF7) from non-CNV REEPIs. Significantly enriched interactions in LTED (adjusted *P* ≤ 0.05; absolute log₂ fold change ≥ 1) include two non-homologous repeat combinations: Low_complexity : LTR/ERVL and Simple_repeat : LTR/ERV1. **b**, *De novo* motif analysis of Low_complexity : LTR/ERVL REEPIs, showing SP1 motifs at promoter anchors and SMAD3 motifs at enhancer anchors. **c**, Motif analysis of Simple_repeat : LTR/ERV1 REEPIs, showing SP1 motifs at promoter anchors and NF-Y (NFYC) motifs at enhancer anchors. **d**, An LTED-enriched Simple_repeat : LTR/ERV1 REEPI links a downstream ERV1-containing enhancer to the *BCAR1* promoter, a gene implicated in endocrine resistance. **e**, Heat map of log₂ fold change in balanced Hi-C contacts between LTED and MCF7 cells (LTED/MCF7; 25-kb bins) across chr16:75,250,000–75,400,000. Both axes represent the same genomic interval. The *BCAR1* promoter and its linked enhancer are indicated, highlighting the strengthened promoter–enhancer contact in LTED cells. **f**, Schematic model proposing that specific transcription factor pairs—SP1–SMAD3 (Low_complexity : LTR/ERVL) and SP1–NFYC (Simple_repeat : LTR/ERV1)—may facilitate enhancer–promoter communication between non-homologous repeat anchors in LTED cells.

Motif analysis of LTED-enriched REEPIs identified two TF pairs with clear anchor specificity. For Low_complexity : LTR/ERVL, SP1 motifs were enriched at promoter repeats, whereas SMAD3 motifs were enriched at enhancer repeats (Fig. 6b; Supplementary Fig. S10). Both TFs were upregulated in LTED (SP1: 5,077 → 8,067; SMAD3: 3,199 → 3,406; normalized RNA-seq (Kato et al., manuscript in preparation). SP1 has been shown to facilitate chromatin looping between enhancers and promoters, with its depletion disrupting defined enhancer–promoter loops^22^. Additionally, SP1–SMAD3 complexes regulate transcription of TGF-β target genes—for example, in pancreatic cells where their interaction is required for TGF-β–induced gene activation^23^. Consistent with these observations, STRING analysis supported a high-confidence protein–protein interaction between SP1 and SMAD3 (Supplementary Fig. 11). For Simple_repeat : LTR/ERV1, SP1 motifs were enriched at the promoter anchors, whereas NF-Y (NFYC) motifs were enriched at enhancer anchors (Fig. 6c; Supplementary Fig. S12). *NFYC* expression increased in LTED (NFYC: 494→ 624 normalized RNA-seq reads (Kato et al., manuscript in preparation)). Previous studies have demonstrated SP1–NFYC cooperativity at GC-rich promoters and in cell-state regulation: Liang et al. showed that SP1 and NF-Y physically cooperate to activate the NPR-A promoter^24^, whereas Nicolás et al. reported co-occupancy of NF-Y and SP-family proteins at the human SP1 gene promoter^25^. Dolfini et al. further reviewed NF-Y’s central role in transcriptional reprogramming and its integration with GC-binding factors such as SP1^26^. Together, these findings support a potential cooperative role for SP1 and NFYC at repeat-rich enhancer–promoter loci, consistent with a high-confidence SP1–NFYC interaction supported by STRING analysis (Supplementary Fig. S13).

To assess whether these TF pairs reflect merely a generic property of enhancer–promoter contacts, we extended motif scanning to all LTED EPIs after excluding those overlapping CNV regions. Motif analysis showed that GC-rich SP1-like motifs were broadly detectable at promoters across nearly all LTED EPIs (Supplementary File 1), consistent with the pervasive GC-rich architecture of mammalian promoters. By contrast, NFYC and SMAD3 motifs were not detectably enriched at enhancer anchors in the CNV-excluded LTED EPI background, yet were preferentially recovered at enhancers of LTED-enriched REEPIs (Supplementary File 2). These motif distributions indicate that the SP1–SMAD3 and SP1–NFYC configurations represent selectively enriched TF pairs associated with a specific subset of REEPIs that is expanded in LTED cells relative to MCF7 cells, rather than a generic consequence of widespread SP1 motif occurrence at promoters.

At the gene level, we further restricted the analysis to LTED-specific REEPIs supported by FIMO motif detection at both promoter and enhancer anchors (*P* < 0.01). This yielded 61 Simple_repeat : LTR/ERV1 and 45 Low_complexity : LTR/ERVL interaction records, corresponding to 92 non-redundant candidate genes (Supplementary Table 1). Representative endocrine-resistance–associated genes included *BCAR1* and *PAK4*, associated with Simple_repeat : LTR/ERV1 REEPIs, and *MALAT1*, associated with Low_complexity : LTR/ERVL REEPIs. Notably, *MALAT1* was also identified in the Simple_repeat : LTR/ERV1 category, suggesting that a single gene locus can participate in multiple repeat-pair configurations in LTED cells.

*BCAR1* encodes a scaffold protein implicated in anti-estrogen resistance, with elevated expression associated with reduced response to tamoxifen and earlier disease recurrence^27^. *PAK4* promotes stemness and disease progression in endocrine-resistant ER-positive breast cancer, whereas its inhibition reverses endocrine-resistant phenotypes^28^. *MALAT1*, a long noncoding RNA that interacts with ER and contributes to ER-regulated transcription, has also been associated with shorter relapse-free survival in tamoxifen-treated ER-positive breast cancer patients^29^. The identification of LTED-specific REEPIs linked to these genes connects repeat-associated enhancer–promoter interactions to pathways previously implicated in adaptation to impaired estrogen signaling and endocrine therapy. A representative Low_complexity : LTR/ERVL REEPI involving the *MALAT1* promoter was detected specifically in LTED cells but not in MCF7 cells (Supplementary Fig. S14a). Motif inspection of the *BCAR1*-associated Simple_repeat : LTR/ERV1 REEPI (Fig. 6d) confirmed SP1-binding motifs within the promoter anchor and NFYC-binding motifs within the paired enhancer anchor, consistent with the SP1–NFYC configuration identified by the global motif analysis (Supplementary Fig. S14b). Consistently, balanced Hi-C contact maps showed increased chromatin contact signals between the *BCAR1* promoter and its linked enhancer region in LTED cells (25-kb resolution; Fig. 6e). A similar increase was observed at the *PAK4* locus (Supplementary Fig. S15), indicating that enhanced chromatin communication is not restricted to a single representative locus.

Collectively, these findings support a model in which specific promoter-repeat : enhancer-repeat pairings provide recruitment platforms for TF combinations that may facilitate enhancer–promoter communication. In particular, SP1–SMAD3 (Low_complexity : ERVL) and SP1–NFYC (Simple_repeat : ERV1) are associated with LTED-enriched REEPIs involving non-homologous repeat anchors. These TF configurations provide a potential explanation for (i) repeat-pair specificity beyond sequence complementarity, (ii) LTED-biased enrichment of selected pairings, and (iii) enhanced chromatin contacts at endocrine resistance-associated gene loci (Fig. 6f). Together, these findings suggest a repeat-encoded regulatory framework associated with transcriptional adaptation and regulatory plasticity during endocrine resistance.

## Discussion

Hormone deprivation imposes profound stress on ER-positive breast cancer cells, necessitating a reprogramming of gene regulatory networks to sustain survival and proliferation. In this study, we systematically mapped CREs and EPIs in estrogen-dependent MCF7 cells and their LTED derivatives, revealing tens of thousands of interactions. Notably, EPIs in both cell states frequently formed complex regulatory networks within TADs, often involving multiple enhancers and promoters. A striking feature of these interactions was their enrichment for REs at both anchor sites, suggesting that REs may substantially contribute to enhancer–promoter communication.

While the genetic and epigenetic underpinnings of endocrine therapy resistance have been extensively characterized, the role of repeat elements in transcriptional adaptation remains largely unexplored^30^. Previous studies have demonstrated that TEs—especially ERVs—can serve as cryptic oncogenic enhancers co-opted by cancer cells^8^. However, their direct involvement in chromatin looping and transcriptional regulation in hormone-resistant breast cancer has been unclear. Our findings demonstrate that REs are not passive genomic bystanders but are actively integrated into enhancer–promoter networks. Specifically, we identified a dynamic remodeling of the repeat landscape in LTED cells: evolutionarily ancient elements, such as older DNA transposons, were slightly depleted at enhancers of REEPIs, whereas younger elements—particularly ERV1 and ERVL LTR retrotransposons—were markedly enriched at LTED-specific REEPIs, indicating selective enrichment of younger repeats during transcriptional reprogramming (Fig. 4). The predominance of ERV1 and ERVL elements at eRNA-producing enhancers in LTED cells suggests a potential role for these repeats in enhancer activation and long-range regulatory interactions associated with transcriptional plasticity under estrogen-deprived conditions^3^.

Functional analysis of REEPI-linked genes further supports the regulatory relevance of these interactions. In LTED cells, REEPIs were preferentially associated with genes involved in chromatin remodeling, DNA repair, stem cell maintenance, and stress adaptation—biological processes relevant to adaptation under endocrine stress. By contrast, MCF7-specific REEPIs were associated with genes linked to developmental and epithelial maintenance programs. Moreover, genes linked to REEPIs were selectively enriched in signaling pathways responsive to hormonal and environmental cues, whereas single-end repeat or non-repeat EPIs were primarily linked to housekeeping processes like RNA processing and translation (Fig. 5; Supplementary Fig. S8). These distinctions link REEPIs to cell state- and stress-associated transcriptional programs, rather than broadly expressed housekeeping functions.

To explore potential mechanisms underlying REEPIs, we examined the sequence composition and TF motif architecture of LTED-enriched REEPIs. Our data support two mechanistic models. First, in a complementary sequence model, homology between the same repeat family at enhancers and promoters could enable direct RNA–RNA duplex formation, tethering loci via base-pairing interactions. Supporting this, approximately 32% of REEPIs featured the same repeat family at both anchors—most frequently Alu, MIR, or simple repeats. These matched pairs may provide a structural scaffold for physical enhancer–promoter proximity. Second, a protein-mediated bridging model posits that distinct repeat families at each anchor may provide recruitment platforms for DNA- or RNA-binding TFs whose interactions could facilitate enhancer–promoter communication. Our motif analyses identified two representative TF pairs: SP1–SMAD3, associated with Low_complexity : ERVL REEPIs, and SP1–NFYC, associated with Simple_repeat : ERV1 REEPIs. Both TF pairs were upregulated in LTED cells, and STRING network analysis supported their high-confidence physical interactions. Prior studies have shown that SP1–SMAD3 and SP1–NFYC can form transcriptionally active complexes, supporting a model in which TFs recognizing distinct repeat-associated motifs may facilitate communication between non-homologous repeat-containing regulatory elements independently of direct sequence homology. Notably, SP1–NFYC–associated REEPIs were linked to key regulators of therapy resistance, including *BCAR1* and *PAK4*, and Hi-C data showed increased enhancer–promoter contacts at both loci in LTED cells. By contrast, SP1–SMAD3-associated REEPIs were linked to therapy-resistance genes such as *MALAT1*. *BCAR1* encodes a scaffold protein whose overexpression confers resistance to anti-estrogen therapies, while *MALAT1* is a lncRNA implicated in metastasis and poor prognosis^27,29^. These examples illustrate how repeat-associated enhancer– promoter interactions may be integrated into regulatory networks involving key mediators of endocrine resistance.

In summary, our study establishes repeat elements as widespread components of the enhancer– promoter interactome in breast cancer cells. A large fraction of EPIs in MCF7 cells (∼86.5%) are associated with REs, and this fraction remains high under hormone-deprived conditions (∼87.8% in LTED). Under estrogen deprivation, LTED cells show selective remodeling of REEPIs involving specific retrotransposon families, particularly ERV1 and ERVL, which are linked to genes involved in chromatin remodeling, stress adaptation, and cell-fate reprogramming. By delineating the sequence features and candidate regulatory mechanisms associated with these interactions, our study reveals how repetitive DNA can be incorporated into regulatory architectures that are remodeled during adaptation to estrogen deprivation. Our findings establish repeat-associated enhancer–promoter communication as a prominent and previously underappreciated feature of the regulatory reprogramming associated with endocrine resistance.

## Methods

### Cell culture and RNA isolation

The human estrogen receptor-positive breast cancer cell line MCF7 (ATCC , Cat# HTB-22) was cultured in RPMI 1640 medium (Thermo Fisher Scientific, Cat# 11875093) supplemented with 10% fetal bovine serum (FBS; Corning, Cat# 35-010-CV). Long-term estrogen-deprived (LTED) cells were established by maintaining MCF7 cells in phenol red-free RPMI 1640 (Thermo Fisher Scientific, Cat# 11835030) supplemented with 4% dextran-coated charcoal-stripped FBS (Thermo Fisher Scientific, Cat# 12676029) for 4 months. All cells were incubated at 37 °C in a humidified atmosphere containing 5% CO₂. Total RNA was extracted from cultured cells using TRIzol reagent (Invitrogen, Cat# 15596018) according to the manufacturer’s instructions.

### CAGE library preparation

Single-strand cap analysis of gene expression (ssCAGE) libraries were prepared following previously published protocols^11,31^. In brief, 1 μg of total RNA was reverse transcribed using random primers. The resulting cDNA was enriched for full-length, capped transcripts via a cap-trapping process. RNase I treatment was then applied to digest uncapped RNA species. The 5′ends of the capped cDNAs were biotinylated and captured using streptavidin-coated magnetic beads. Following linker ligation and PCR amplification, libraries were purified and sequenced using an Illumina HiSeq 2500 platform in high-output mode. CAGE tags were aligned to the human reference genome to identify transcription start sites (TSSs) at single-nucleotide resolution.

### CAGE analysis and annotation of promoters and enhancers

Promoters and enhancers were identified using the SCAFE pipeline^13^, a computational framework designed to detect cis-regulatory elements (CREs) from Cap Analysis of Gene Expression (CAGE) data. SCAFE was run with default parameters, which merge transcription start site (TSS) clusters located within +100 nucleotides (nt) downstream and –400 nt upstream into a single CRE unit. CREs were annotated using gene models from GENCODE release 38 based on the GRCh38/hg38 genome assembly. Regulatory elements located within ±500 nt of an annotated TSS on the same strand were classified as promoters, while all remaining CREs were designated as enhancers.

### RADICL-seq mapping and processing

RADICL-seq raw sequencing data were obtained from the DNA Data Bank of Japan (DDBJ) under accession number DRA19579. Mapping and downstream processing were performed following previously published protocols^32^ (Kato et al., manuscript in preparation). Briefly, chimeric RNA–DNA reads were aligned to the GRCh38/hg38 genome assembly using standard RADICL-seq alignment pipelines. The statistical significance of RNA–DNA interactions was calculated as previously described, and interactions with an adjusted *P* (q-value) ≤ 0.001 were retained for downstream analysis.

### High-confidence interaction filtering and enhancer–promoter interaction identification

To ensure robust detection of enhancer–promoter interactions (EPIs), we applied a background correction step using a modified negative binomial model implemented in the CHiCANE R package (v0.1.8)^33^. This statistical framework accounts for biases introduced by genomic distance and contact frequency. Only RNA–DNA interaction pairs with an adjusted *P* (q-value) ≤ 0.001 were retained for downstream analysis. EPIs were identified by integrating the filtered RADICL-seq interactions with CAGE-defined cis-regulatory elements (CREs), assuming that promoter-derived RNAs and enhancer RNAs remain associated with their sites of transcription while interacting with their cognate regulatory elements. A two-step assignment strategy was implemented using BEDTools (v2.26.0)^34^ to annotate interactions based on the genomic origin of RNA tags and the target loci of DNA tags. In Rule 1, RNA tags aligning to promoter regions on the same strand were paired with DNA tags overlapping enhancer regions (pRNA–eDNA). In Rule 2, RNA tags aligning to enhancer regions—regardless of strand orientation—were paired with DNA tags overlapping promoter regions (eRNA–pDNA). To minimize potential artifacts, interactions in which both anchors mapped to the same CRE (i.e., CRE self-interactions) were excluded from the analysis.

### EPI–TAD boundary enrichment analysis

To examine the spatial distribution of EPIs relative to topologically associating domains (TADs), we conducted a genome-wide meta-profile analysis of EPI anchor positions. Each TAD was symmetrically extended upstream and downstream by one TAD length, resulting in a region encompassing 3× the original TAD size. This extended region was divided into 300 equal bins: 100 bins upstream, 100 bins within the TAD body, and 100 bins downstream. The midpoint of each EPI anchor was calculated and mapped to its corresponding bin within the normalized TAD window. Anchor positions were aggregated across all TADs, and the average density per bin was computed. The resulting signal was smoothed using a 3-bin rolling average and Z-score normalized across bins. TAD boundaries were defined as ±10% around the start (0%) and end (100%) positions of the normalized window. All other bins were designated as “Other” regions. Average Z-scores in boundary bins were compared to those in “Other” regions using a two-tailed Student’s t-test.

### Motif analysis of enhancers and promoters involved in enhancer–promoter interactions

De novo motif discovery was performed separately on enhancer and promoter sequences derived from EPIs. STREME (MEME Suite v5.5.8)^35^ was used with the following parameters: --minw 4 --maxw 30 --thresh 0.05. A shuffled version of the input sequences served as a background control. To assess whether EPI-associated motifs showed significant clustering within the AluSx consensus sequence, STREME-derived motifs were mapped to the AluSx consensus sequence using MCAST (MEME Suite v5.5.8)^36^.

### Definition and identification of REEPIs

The repeat element annotations were obtained from RepeatMasker (v4.1.5) (https://www.repeatmasker.org/). To identify REEPIs, enhancer and promoter coordinates from the EPIs were intersected with repeat element annotations using BEDTools (v2.26.0)^34^. EPIs in which repeat elements overlapped both the enhancer and promoter anchors were defined as REEPIs.

### Differential enrichment analysis of repeat element families

To assess the enrichment of repeat element families in EPI anchors, we quantified the number of enhancer and promoter anchors—separately in MCF7 and LTED cells—that overlapped annotated repeat families. Both single-end repeat EPIs and REEPIs were included in the analysis. Interactions involving a single enhancer paired with multiple promoters, or a single promoter paired with multiple enhancers, were treated as distinct EPIs. As individual CREs can overlap with multiple repeat elements, each overlapping repeat was counted independently when quantifying repeat composition within REEPIs. However, the total number of EPIs was treated as a set of discrete interaction events and was not weighted by repeat multiplicity. Differential enrichment between cell types was determined using the edgeR package (v3.40.2)^37^. Repeat families with adjusted *P* ≤ 0.05 and an absolute log₂ fold change ≥ 0.585 (1.5-fold) were considered significantly enriched.

### Gene Ontology (GO) analysis

Gene symbols were converted to Entrez Gene IDs using the *bitr* function from the clusterProfiler R package (v4.6.2)^38^. GO enrichment analysis for REEPI-associated genes was performed using the *enrichGO* function with default parameters, focusing on the Biological Process (BP) category.

### Differential interaction analysis of repeat element pairs

Repeat–repeat interaction counts were normalized using the Trimmed Mean of M-values (TMM) method implemented in edgeR (v3.40.2)^37^. Differential interaction analysis between MCF7 and LTED cells was then performed using the same package. Repeat pairs with *adjusted P* ≤ 0.05 and an |log₂ fold change| ≥ 1 were considered significantly altered. Volcano plots were generated with ggplot2 (v3.4.2)^39^ to visualize differentially enriched repeat–repeat interactions.

### Motif analysis of differentially enriched repeat–repeat pairs in LTED

To examine potential sequence-specific features of repeat–repeat interactions enriched in LTED cells, we performed separate *de novo* motif analyses for repeat elements located at enhancer and promoter anchors. STREME (MEME Suite v5.5.8)^35^ was run with parameters --minw 6 --maxw 15 --thresh 0.05. Identified motifs were compared against known transcription factor binding sites in the HOCOMOCO v12 human core database using Tomtom (MEME Suite v5.5.8)^40^ under default settings. For motif inspection, we used FIMO (MEME Suite v5.5.8)^41^ to identify input sequences containing the selected motifs.

### Hi-C contact analysis

Local chromatin contacts between MCF7 and LTED cells at the *BCAR1* locus were compared using balanced 25-kb Hi-C contact maps extracted from chr16:75,250,000–75,400,000 with cooler (v0.10.4)^42^ on the MCF7 and LTED .cool files (Kato et al., manuscript in preparation). Log₂ fold-change values (LTED / MCF7) were calculated for each 25-kb bin pair after merging the two matrices in R and filling missing values with zeroes. These log₂ fold-change values were then visualized as a heat map using the ggplot2 (ver. 3.4.2)^39^ R package, highlighting increased LTED contacts between the *BCAR1* promoter and its linked enhancer.

### Visualization

The plots of this research were performed by IGV Web App (https://igv.org/app/) and ggplot2 (ver. 3.4.2)^39^ R package.

## Data availability

All sequencing data generated in this study have been deposited in the Gene Expression Omnibus (GEO) and the DDBJ Sequence Read Archive (DRA). The CAGE datasets are available under accession number GSE299910. The RADICL-seq data are available under accession number DRA19579. The RNA-seq and Hi-C datasets, including processed TAD annotations, are available under accession numbers DRA009612 and DRA026723, respectively.

## Acknowledgments

This work was supported by the RIKEN Center for Integrative Medical Sciences under the auspices of the Japanese Ministry of Education, Culture, Sports, Science and Technology (MEXT); JSPS KAKENHI Grant Numbers JP19K06623, JP22K06187, and JP25K09598 (to M.K.), JP23H00411, and JP24H01382 (to N.S.), and JP25K18864 (to M.P.); grants from the Japanese Cancer Association and the Kobayashi Foundation for Cancer Research (JCA-KFCR) (to M.P.); and grants from the Takeda Science Foundation and the Naito Foundation (to N.S.); RIKEN BDR intramural grants; the RIKEN Pioneering Project “Genome Building from TADs”; JST CREST Grant Number JPMJCR20S5; JSPS KAKENHI Grant Numbers JP20K20582 and JP25H00982 (I.H.) and JSPS KAKENHI Grant Number JP25K24823 (to H.M.).

This work was also supported by the Human Technopole Foundation, funded by the Italian Government through the Ministries of Economy and Finance, Health, and University and Research. This publication was also carried out with co-financing from the European Union – Next Generation EU, MISSION 4, COMPONENT 2, ‘From Research to Business’, INVESTMENT 1.4, ‘Strengthening research infrastructures and creation of national R&D champions’, in relation to the project identified by code CN00000041 _S6_LA_006, titled ‘National Center for Gene Therapy and Drugs based on RNA Technology and CUP CNR B83C22002860006’.

Sequencing was performed by the Laboratory for Genotyping Development at RIKEN IMS. We thank Nobuyuki Takeda and Teruaki Kitakura (RIKEN IMS) for IT infrastructure support for the FANTOM6 collaboration, Emi Ito (RIKEN IMS) for administrative support and Akie Tanigawa (RIKEN BDR) for technical assistance with Hi-C library preparation. We also thank all members of the FANTOM6 consortium for valuable discussions and suggestions.

## Author contributions

X.S. and M.K. conceived and designed the study. X.S. and S.N. performed the bioinformatic analyses.

M.P. performed the wet-lab experiments. H.M. and I.H. contributed the Hi-C datasets. M.K., N.S., and P.C. supervised the study. X.S., S.N., and M.K. wrote the manuscript with input from all authors. C.Y., H.T., T.K., M.K., and P.C. coordinated the datasets as part of the FANTOM6 headquarters.

## References

1. Lander, E.S. et al. Initial sequencing and analysis of the human genome. Nature 409, 860–921 (2001).

2. Feschotte, C. Transposable elements and the evolution of regulatory networks. Nat Rev Genet 9, 397–405 (2008).

3. Chuong, E.B., Elde, N.C. & Feschotte, C. Regulatory activities of transposable elements: from conflicts to benefits. Nat Rev Genet 18, 71–86 (2017).

4. Sundaram, V. et al. Functional cis-regulatory modules encoded by mouse-specific endogenous retrovirus. Nat Commun 8, 14550 (2017).

5. Wicker, T. et al. A unified classification system for eukaryotic transposable elements. Nat Rev Genet 8, 973–82 (2007).

6. Rebollo, R., Romanish, M.T. & Mager, D.L. Transposable elements: an abundant and natural source of regulatory sequences for host genes. Annu Rev Genet 46, 21–42 (2012).

7. Jang, H.S. et al. Transposable elements drive widespread expression of oncogenes in human cancers. Nat Genet 51, 611–617 (2019).

8. Babaian, A. & Mager, D.L. Endogenous retroviral promoter exaptation in human cancer. Mob DNA 7, 24 (2016).

9. Osborne, C.K. & Schiff, R. Mechanisms of endocrine resistance in breast cancer. Annu Rev Med 62, 233–47 (2011).

10. Tomita, S. et al. A cluster of noncoding RNAs activates the ESR1 locus during breast cancer adaptation. Nat Commun 6, 6966 (2015).

11. Takahashi, H., Lassmann, T., Murata, M. & Carninci, P. 5’ end-centered expression profiling using cap-analysis gene expression and next-generation sequencing. Nat Protoc 7, 542–61 (2012).

12. Bonetti, A. et al. RADICL-seq identifies general and cell type-specific principles of genome-wide RNA-chromatin interactions. Nat Commun 11, 1018 (2020).

13. Moody, J. et al. SCAFE: a software suite for analysis of transcribed cis-regulatory elements in single cells. Bioinformatics 38, 5126–5128 (2022).

14. Liang, L. et al. Complementary Alu sequences mediate enhancer-promoter selectivity. Nature 619, 868–875 (2023).

15. Rao, S.S. et al. A 3D map of the human genome at kilobase resolution reveals principles of chromatin looping. Cell 159, 1665–80 (2014).

16. Akgol Oksuz, B., et al. Systematic evaluation of chromosome conformation capture assays. Nat Methods 18, 1046–1055 (2021).

17. Schmitt, A.D., Hu, M. & Ren, B. Genome-wide mapping and analysis of chromosome architecture. Nat Rev Mol Cell Biol 17, 743–755 (2016).

18. Dixon, J.R. et al. Topological domains in mammalian genomes identified by analysis of chromatin interactions. Nature 485, 376–80 (2012).

19. Robberecht, C., Voet, T., Zamani Esteki, M., Nowakowska, B.A. & Vermeesch, J.R. Nonallelic homologous recombination between retrotransposable elements is a driver of de novo unbalanced translocations. Genome Res 23, 411–8 (2013).

20. Cardoso, A.R., Oliveira, M., Amorim, A. & Azevedo, L. Major influence of repetitive elements on disease-associated copy number variants (CNVs). Hum Genomics 10, 30 (2016).

21. Chen, L., Zhou, W., Zhang, L. & Zhang, F. Genome architecture and its roles in human copy number variation. Genomics Inform 12, 136–44 (2014).

22. Deshane, J. et al. Sp1 regulates chromatin looping between an intronic enhancer and distal promoter of the human heme oxygenase-1 gene in renal cells. J Biol Chem 285, 16476–86 (2010).

23. Jungert, K. et al. Smad-Sp1 complexes mediate TGFbeta-induced early transcription of oncogenic Smad7 in pancreatic cancer cells. Carcinogenesis 27, 2392–401 (2006).

24. Liang, F., Schaufele, F. & Gardner, D.G. Functional interaction of NF-Y and Sp1 is required for type a natriuretic peptide receptor gene transcription. J Biol Chem 276, 1516–22 (2001).

25. Nicolás, M., Noé, V. & Ciudad, C.J. Transcriptional regulation of the human Sp1 gene promoter by the specificity protein (Sp) family members nuclear factor Y (NF-Y) and E2F. Biochem J 371, 265–75 (2003).

26. Dolfini, D., Imbriano, C. & Mantovani, R. The role(s) of NF-Y in development and differentiation. Cell Death Differ 32, 195–206 (2025).

27. van der Flier, S. et al. Bcar1/p130Cas protein and primary breast cancer: prognosis and response to tamoxifen treatment. J Natl Cancer Inst 92, 120–7 (2000).

28. Santiago-Gómez, A. et al. PAK4 regulates stemness and progression in endocrine resistant ER-positive metastatic breast cancer. Cancer Lett 458, 66–75 (2019).

29. Huang, N.S. et al. Long non-coding RNA metastasis associated in lung adenocarcinoma transcript 1 (MALAT1) interacts with estrogen receptor and predicted poor survival in breast cancer. Oncotarget 7, 37957–37965 (2016).

30. Jeselsohn, R., Buchwalter, G., De Angelis, C., Brown, M. & Schiff, R. ESR1 mutations—a mechanism for acquired endocrine resistance in breast cancer. Nat Rev Clin Oncol 12, 573–83 (2015).

31. Morioka, M.S. et al. Cap Analysis of Gene Expression (CAGE): A Quantitative and Genome-Wide Assay of Transcription Start Sites. Methods Mol Biol 2120, 277–301 (2020).

32. Shu, X., Kato, M., Takizawa, S., Suzuki, Y. & Carninci, P. RADIP technology comprehensively identifies H3K27me3-associated RNA-chromatin interactions. Nucleic Acids Res 52, e104 (2024).

33. Holgersen, E.M. et al. Identifying high-confidence capture Hi-C interactions using CHiCANE. Nat Protoc 16, 2257–2285 (2021).

34. Quinlan, A.R. & Hall, I.M. BEDTools: a flexible suite of utilities for comparing genomic features. Bioinformatics 26, 841–2 (2010).

35. Bailey, T.L. STREME: accurate and versatile sequence motif discovery. Bioinformatics 37, 2834–2840 (2021).

36. Bailey, T.L. & Noble, W.S. Searching for statistically significant regulatory modules. Bioinformatics 19 Suppl 2, ii16-25 (2003).

37. Robinson, M.D., McCarthy, D.J. & Smyth, G.K. edgeR: a Bioconductor package for differential expression analysis of digital gene expression data. Bioinformatics 26, 139–40 (2010).

38. Wu, T. et al. clusterProfiler 4.0: A universal enrichment tool for interpreting omics data. Innovation (Camb) 2, 100141 (2021).

39. Wickham, H. Ggplot2 : Elegant Graphics for Data Analysis., (Springer-Verlag New York, 2016).

40. Gupta, S., Stamatoyannopoulos, J.A., Bailey, T.L. & Noble, W.S. Quantifying similarity between motifs. Genome Biol 8, R24 (2007).

41. Grant, C.E., Bailey, T.L. & Noble, W.S. FIMO: scanning for occurrences of a given motif. Bioinformatics 27, 1017–8 (2011).

42. Abdennur, N. & Mirny, L.A. Cooler: scalable storage for Hi-C data and other genomically labeled arrays. Bioinformatics 36, 311–316 (2020).

