## Supplementary material for "Repeat element–anchored enhancer–promoter interactions (REEPIs) shape the regulatory landscape of hormone-resistant breast cancer": Sup Figs

Affiliations:
1. RIKEN Center for Integrative Medical Sciences, Yokohama Kanagawa 230-0045, Japan.
2. The Department of Computational Biology and Medical Sciences, The Graduate School of Frontier Sciences, University of Tokyo, 5-1-5 Kashiwanoha, Kashiwa, Chiba 277-8561, Japan.
3. Division of Cancer Biology, The Cancer Institute of JFCR, 3-8-31 Ariake, Koto-ku, Tokyo 135-8550, Japan.
4. Laboratory for Developmental Epigenetics, RIKEN Center for Biosystems Dynamics Research (BDR), Kobe, Japan
5. Human Technopole, Milan, 20157, Italy
6. Medical Institute of Bioregulation, Kyushu University
7. These authors contributed equally: Xufeng Shu, Shengliang Ni
8. Present address: School of Life Science and Technology, Institute of Science Tokyo, 2-12-1-M6-12, Ookayama, Meguro-ku, Tokyo, 152-8550, Japan

∗

- Supplementary Figures S1- 15 (PDF)

- Supplementary File 1_promoter (STREAM Results) (PDF)

- Supplementary File 2_enhancer (STREAM Results) (PDF)

- Supplementary Table 1 (excel file)

#### Supplementary Figures

a

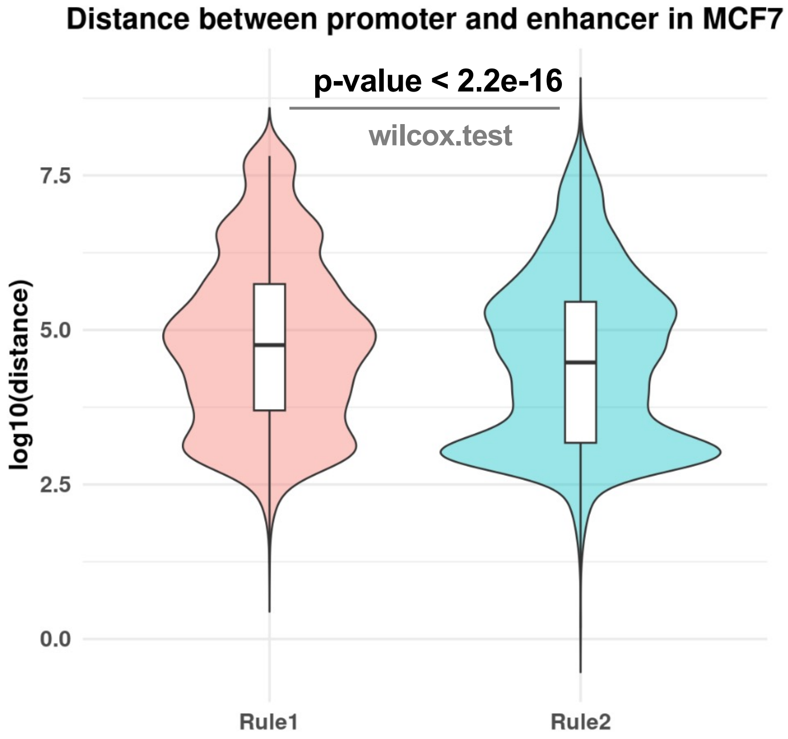


b

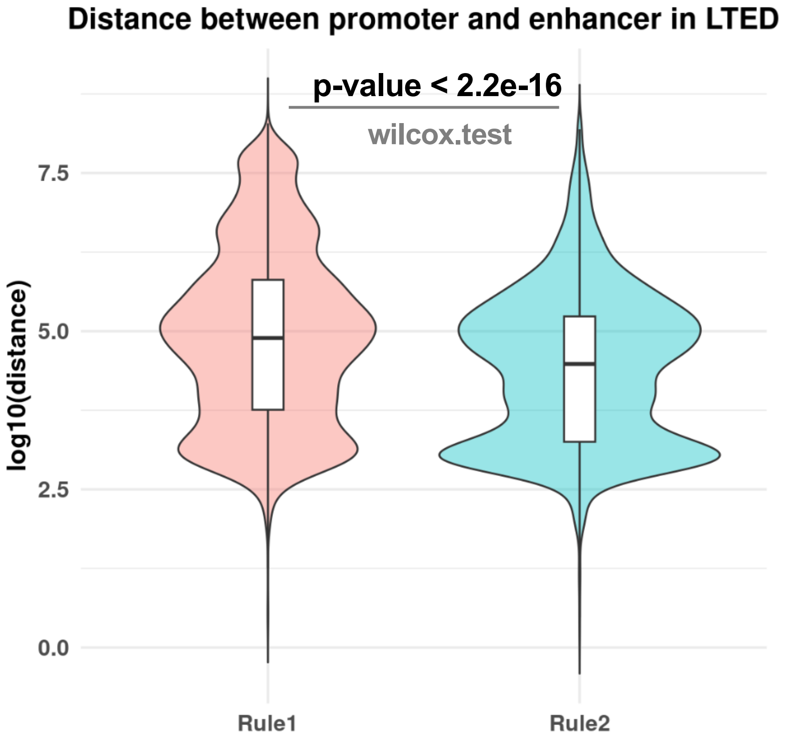


**Supplementary Figure S1: Genomic distance between promoter and enhancer in Rule 1 and Rule 2 EPIs. a**, Violin plot showing the distribution of genomic distances between promoter and enhancer anchors for Rule 1 and Rule 2 EPIs in MCF7 cells. Rule 1 EPIs span significantly greater distances than Rule 2 EPIs. **b**, Same analysis as in (a) for LTED cells, showing a consistent trend of longer genomic distances for Rule 1 EPIs.

#### Supplementary Figure S1

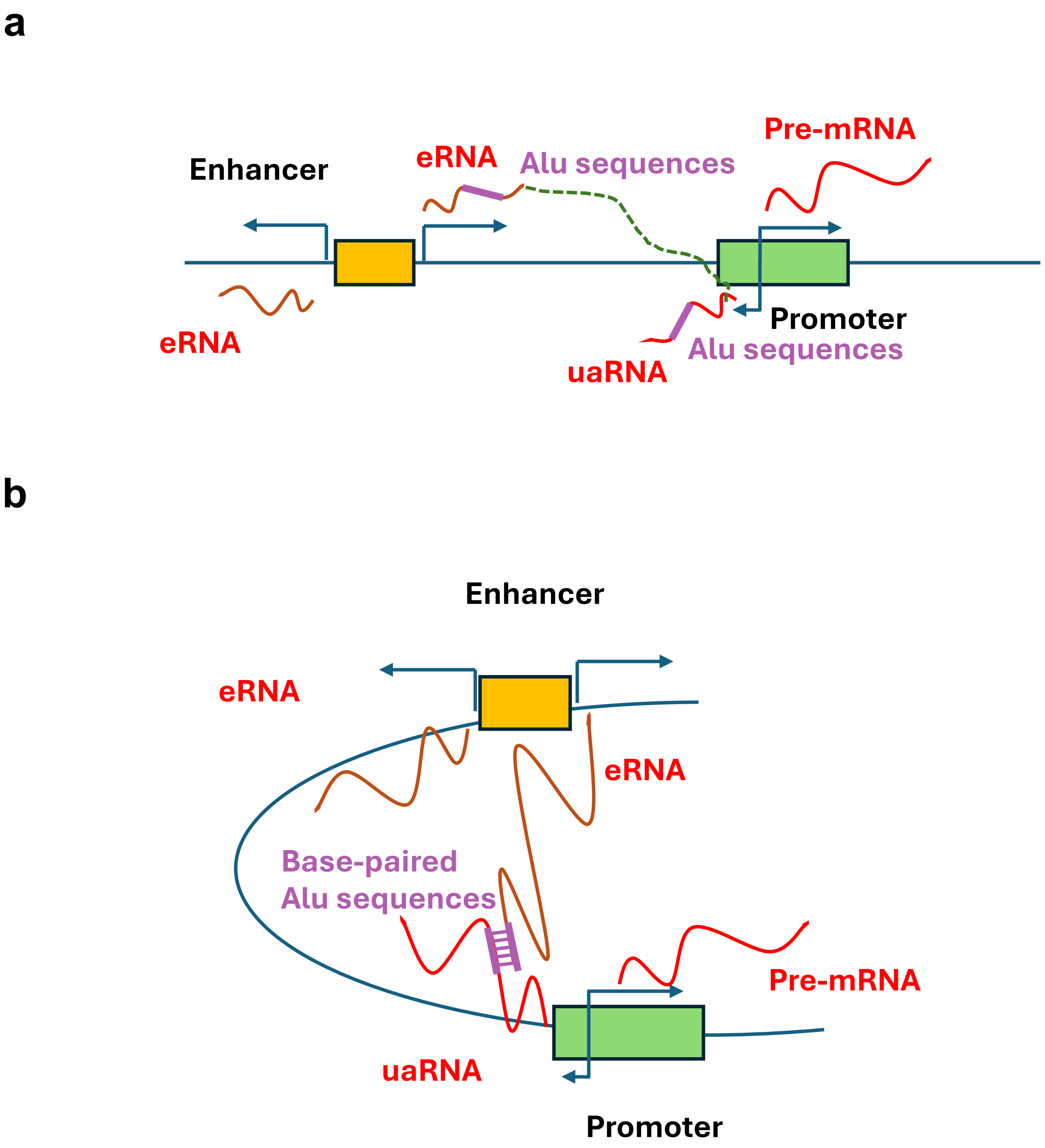


**Supplementary Figure S2: RNA–RNA pairing via Alu elements mediates enhancer–promoter specificity. a**, Model depicting enhancer–promoter interactions mediated by base-pairing between eRNAs and uaRNAs (PROMPTs) transcribed from promoters. **b**, Schematic showing chromatin looping stabilized by complementary Alu sequences embedded in both eRNAs and uaRNAs.

#### Supplementary Figure S2

**a**

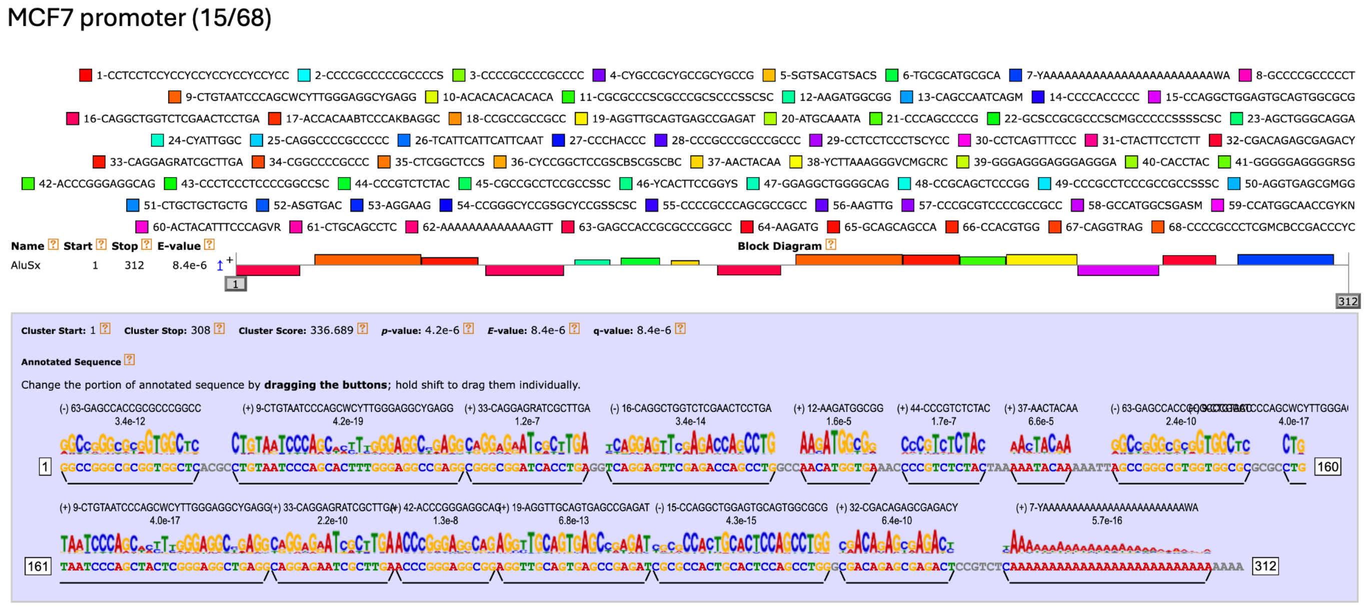


**b**

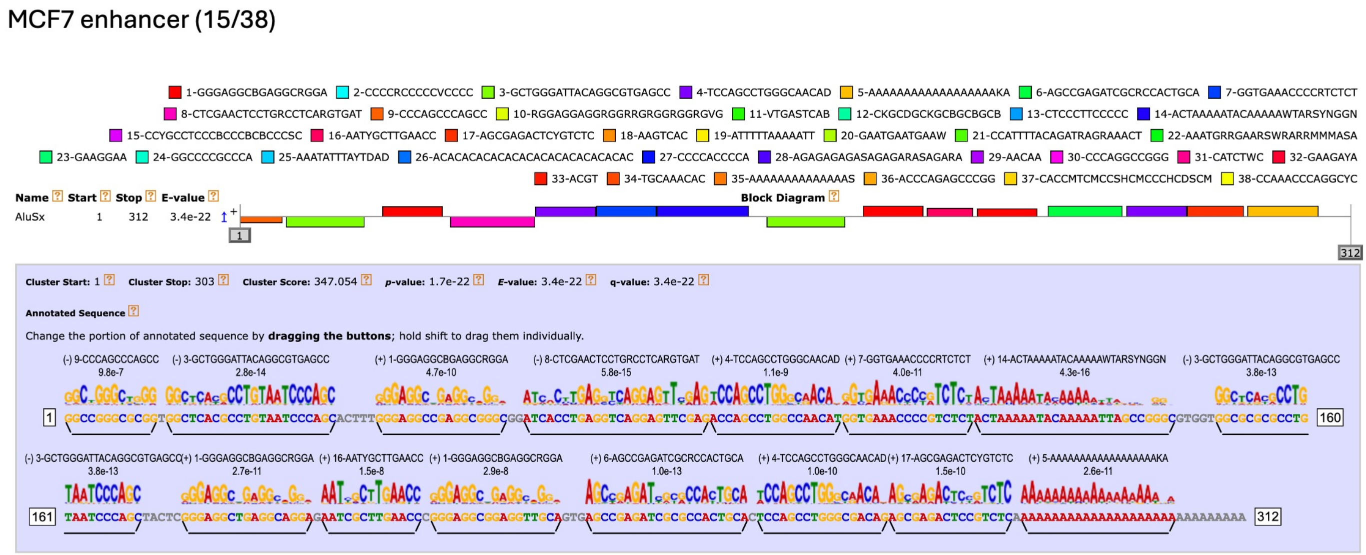


**Supplementary Figure S3: EPI-associated motifs show significant clustering within the AluSx consensus sequence in MCF7 cells. a,** De novo motifs identified from promoter regions involved in EPIs were mapped to the AluSx consensus sequence using MCAST, revealing significant motif clustering ($P = 4.2 \times 10^{-6}$). **b,** Motifs identified from EPI-associated enhancer regions similarly showed significant clustering within the AluSx consensus sequence ($P = 1.7 \times 10^{-22}$), consistent with a shared Alu-associated sequence signature at EPI anchors.

### Supplementary Figure S3

**a**

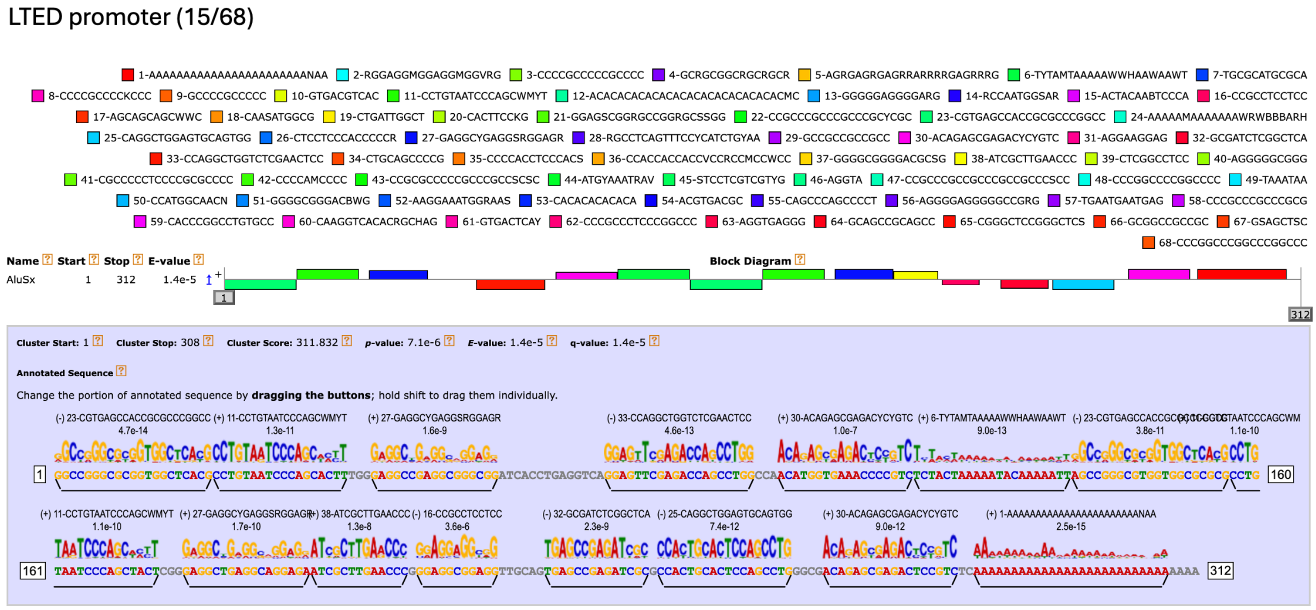


**b**

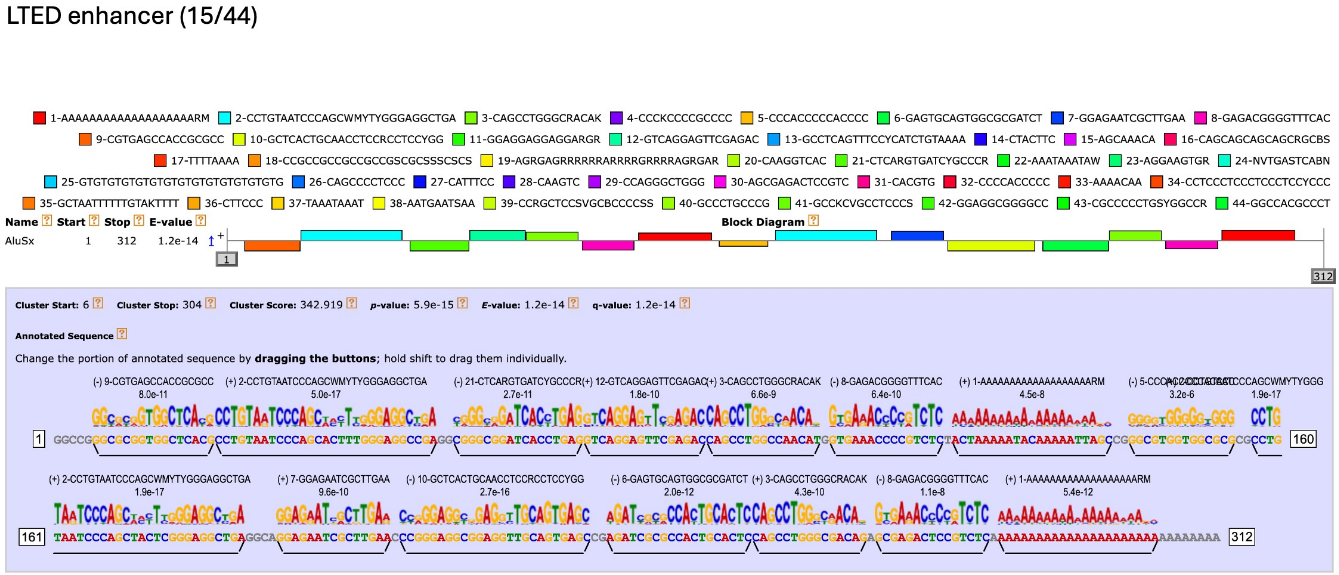


**Supplementary Figure S4: EPI-associated motifs show significant clustering within the AluSx consensus sequence in LTED cells. a,** De novo motifs identified from promoter regions involved in EPIs were mapped to the AluSx consensus sequence using MCAST, revealing significant motif clustering ($P = 7.1 \times 10^{-6}$). **b,** Motifs identified from EPI-associated enhancer regions similarly showed significant clustering within the AluSx consensus sequence ($P = 5.9 \times 10^{-15}$), consistent with a shared Alu-associated sequence signature at EPI anchors.

### Supplementary Figure S4

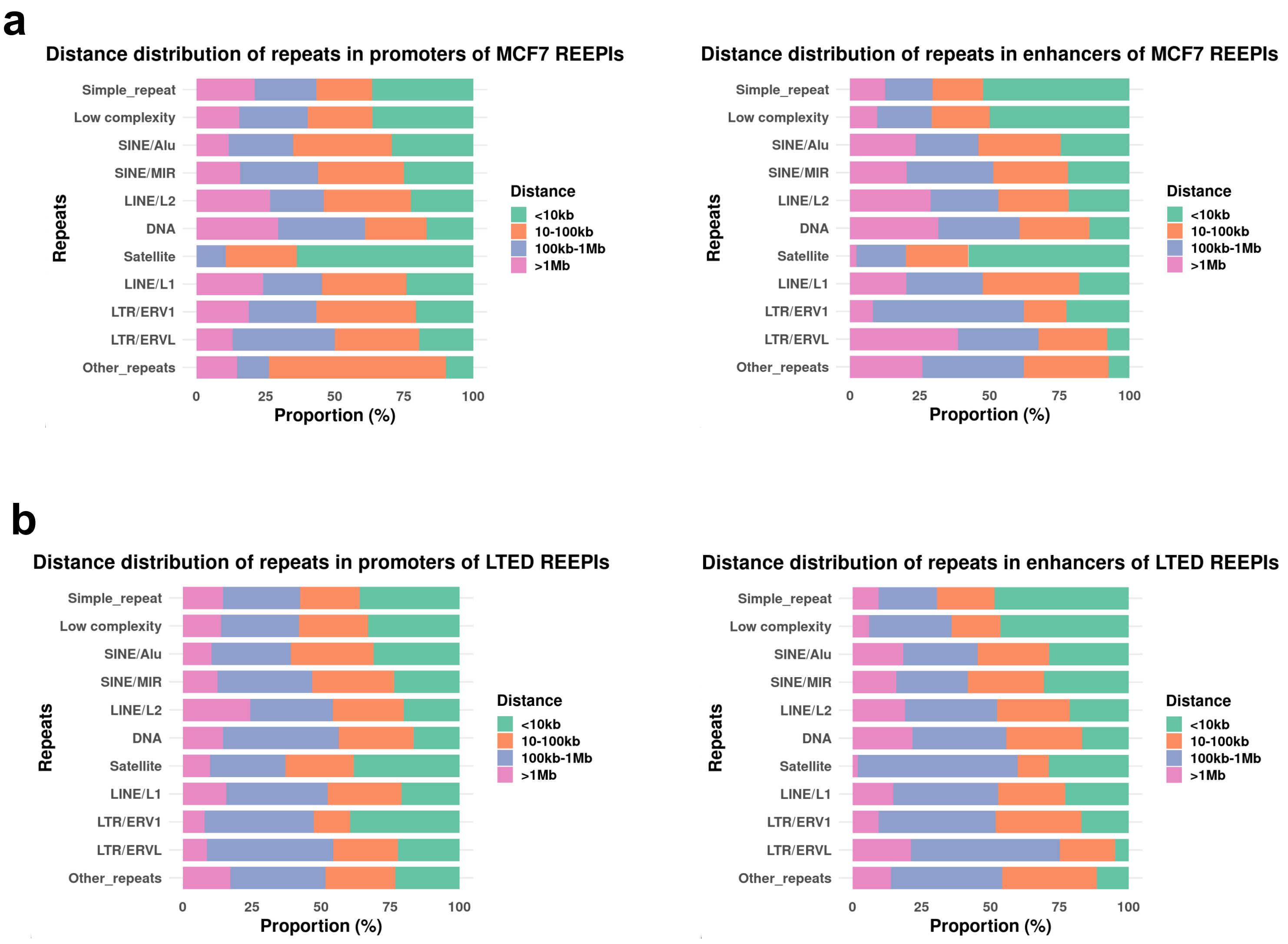


**Supplementary Figure S5: Genomic distance–dependent distribution of repeat subclasses at enhancer–promoter interaction anchors. a**, Distribution of repeat subclasses in promoter (left) and enhancer (right) regions of REEPIs in MCF7 cells, stratified by interaction distance: <10 kb, 10–100 kb, 100 kb–1 Mb, and >1 Mb. **b**, Corresponding distributions for REEPIs in LTED cells.

### Supplementary Figure S5

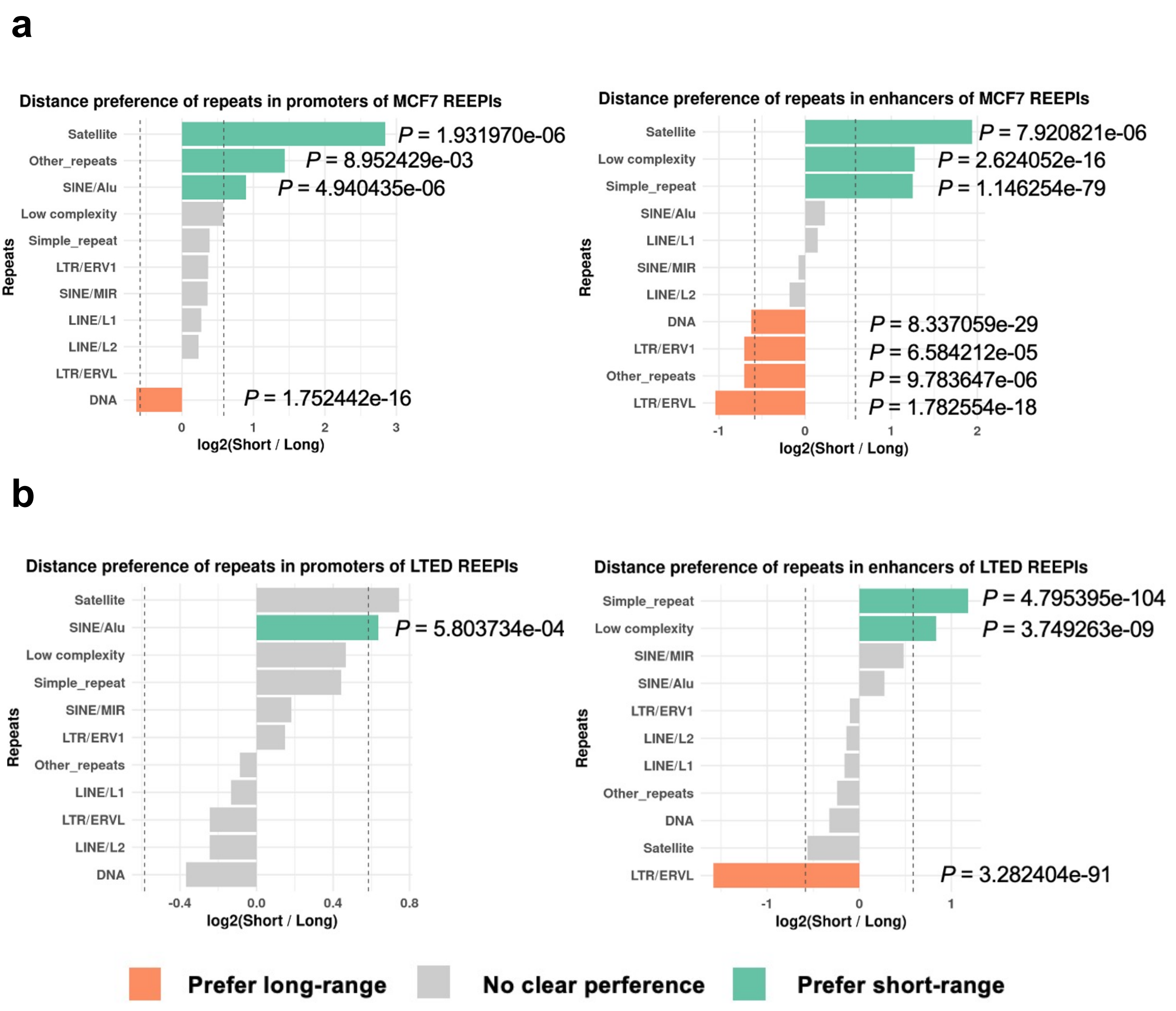


**Supplementary Figure S6: Distance preference of repeat subclasses at enhancer–promoter interaction anchors. a**, Distance preference of repeat subclasses in the promoter (left) and enhancer (right) regions of MCF7 REEPIs. **b**, Corresponding analyses for LTED REEPIs. Bars represent $\log_2$(Short/Long), where Short = (<10 kb + 10–100 kb) and Long = (100 kb–1 Mb + >1 Mb). Statistical significance was determined by two-sided Fisher's exact test (*P* ≤ 0.05; |$\log_2$ fold change| ≥ 0.585, corresponding to ≥ 1.5-fold difference).

#### Supplementary Figure S6

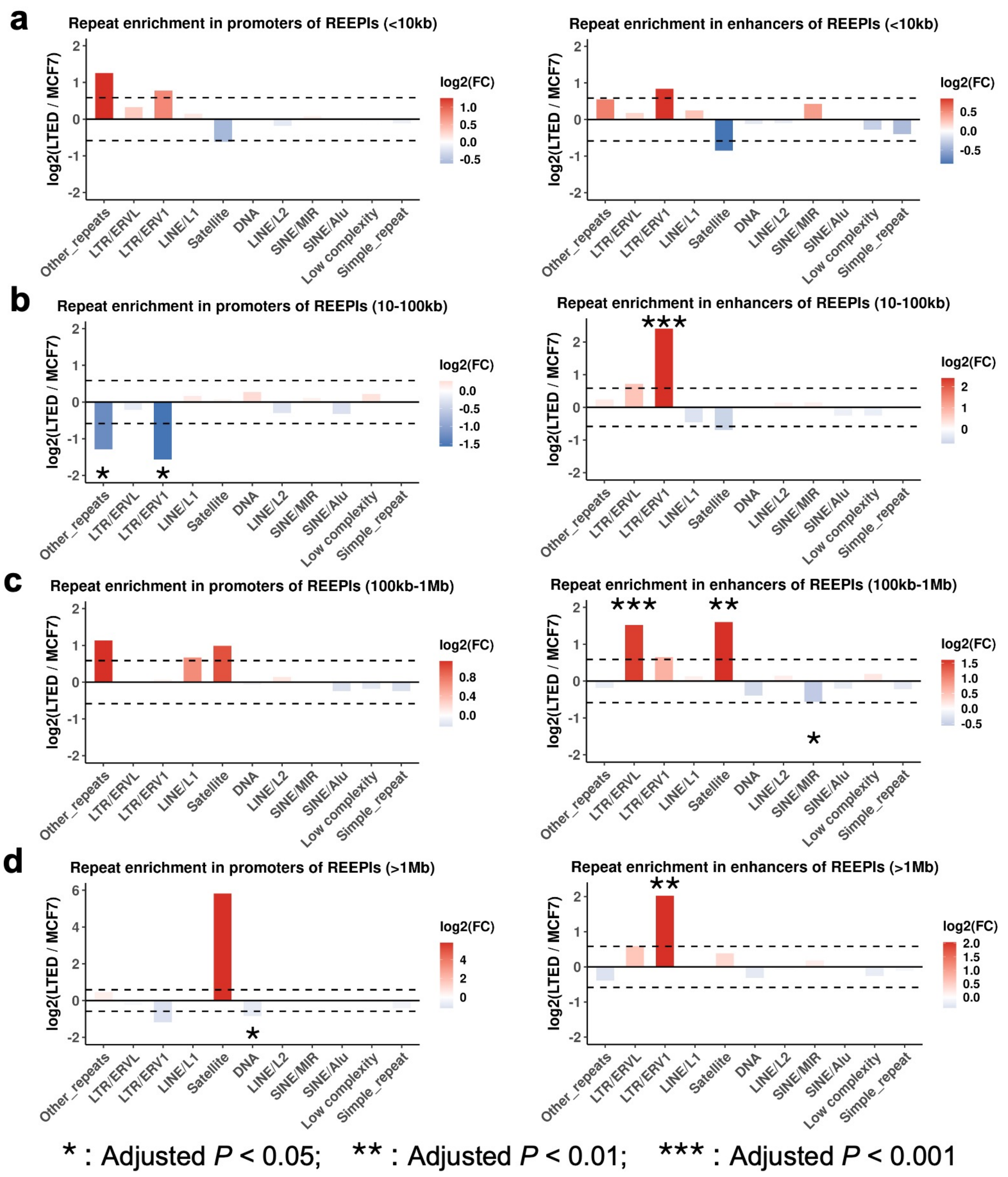


**Supplementary Figure S7: Distance-resolved enrichment of repeat subclasses in REEPIs of LTED relative to MCF7 cells. a–d,** Enrichment of repeat subclasses in the promoter (left) and enhancer (right) anchors of REEPIs across four genomic distance ranges: < 10 kb (a), 10–100 kb (b), 100 kb–1 Mb (c), and > 1 Mb (d). Statistical significance was determined by two-sided Fisher's exact test (P ≤ 0.05; |$\log_2$ fold change| ≥ 0.585, corresponding to ≥ 1.5-fold difference).

### Supplementary Figure S7

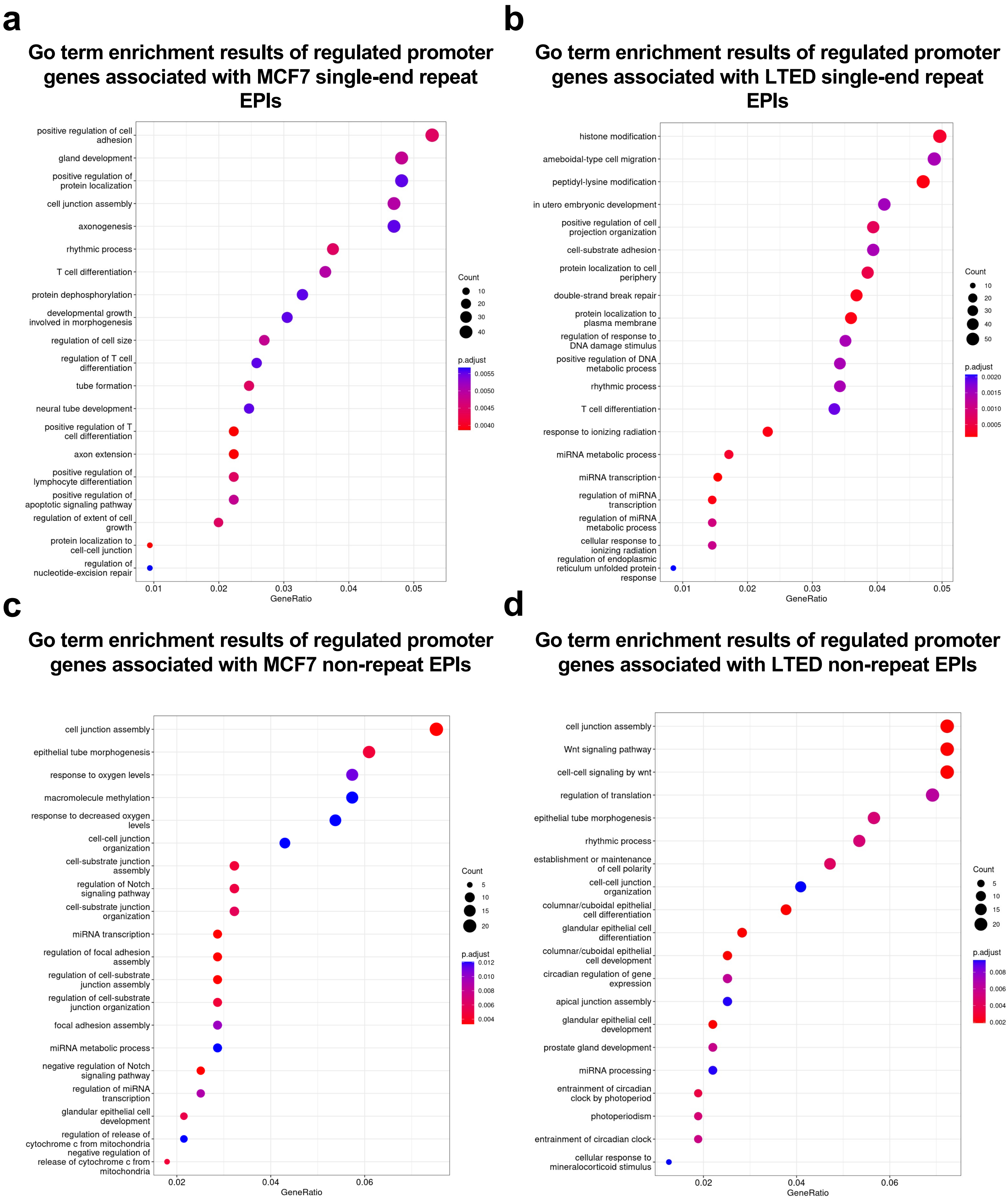


**Supplementary Figure S8: GO enrichment for genes linked to single-end and non-repeat EPIs. a**, MCF7 single-end repeat EPIs. **b**, LTED single-end repeat EPIs. **c**, MCF7 non-repeat EPIs. **d**, LTED non-repeat EPIs.

### Supplementary Figure S8

a

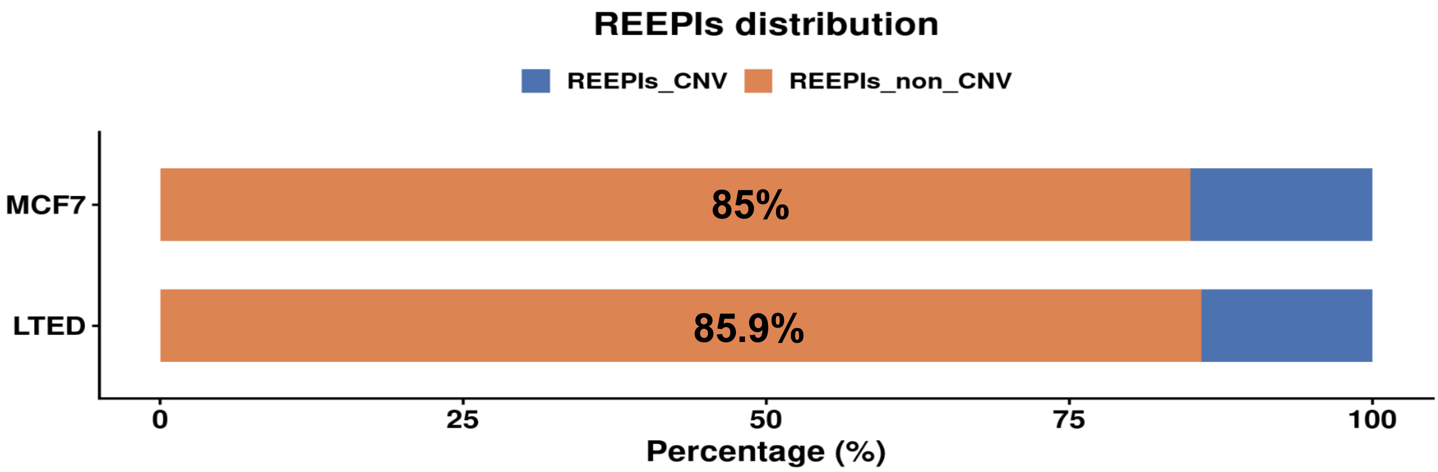


b

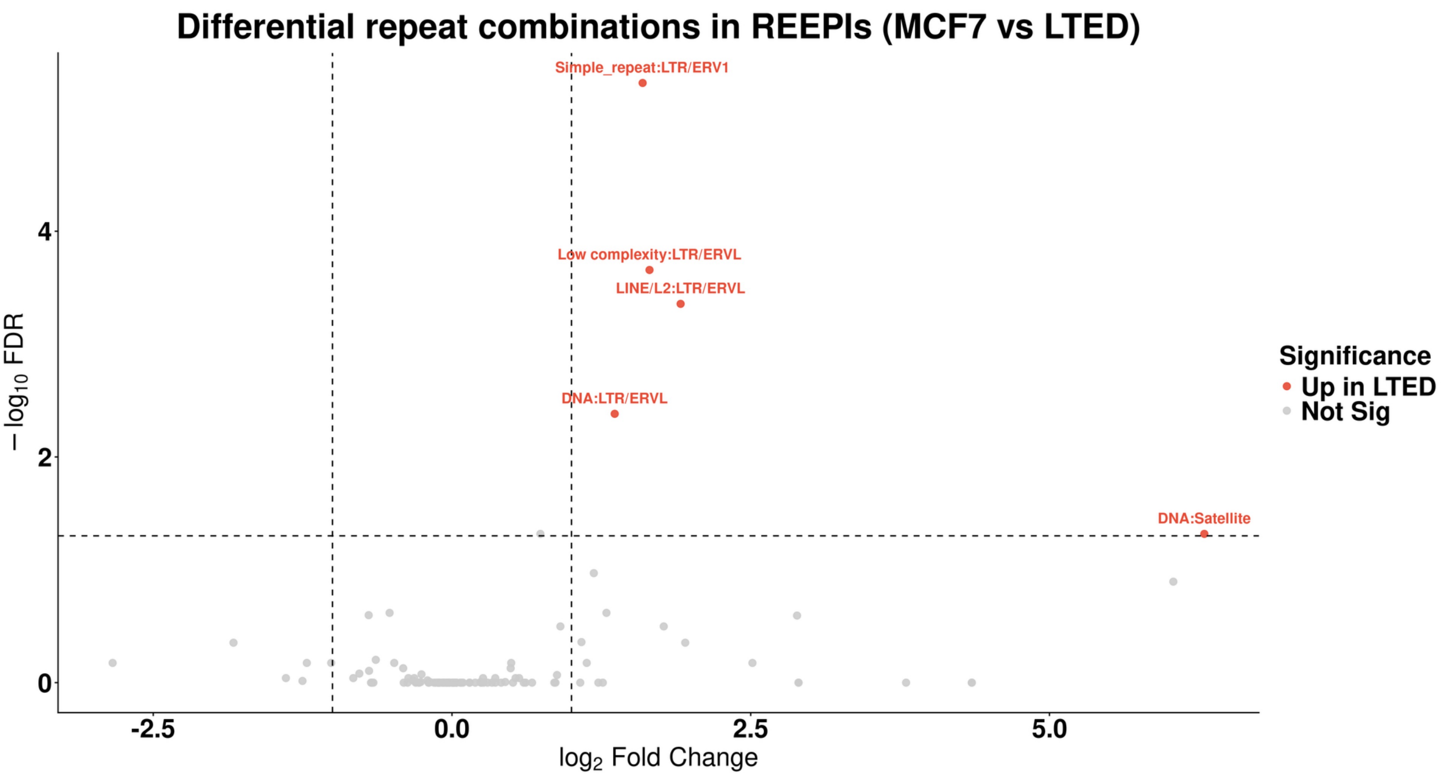


**Supplementary Figure S9: Differential enrichment of repeat–repeat interaction pairs between LTED and MCF7 cells based on CNV-included REEPIs. a**, Bar plot showing the percentages of REEPIs located outside of CNV regions in MCF7 and LTED cells. Most REEPIs lie outside CNV regions. **b**, Volcano plot showing significantly enriched repeat–repeat interaction pairs in LTED versus MCF7 cells (adjusted $P \leq 0.05$; $|\log_2$ fold change$| \geq 1$), based on REEPIs including CNV-overlapping regions.

### Supplementary Figure S9

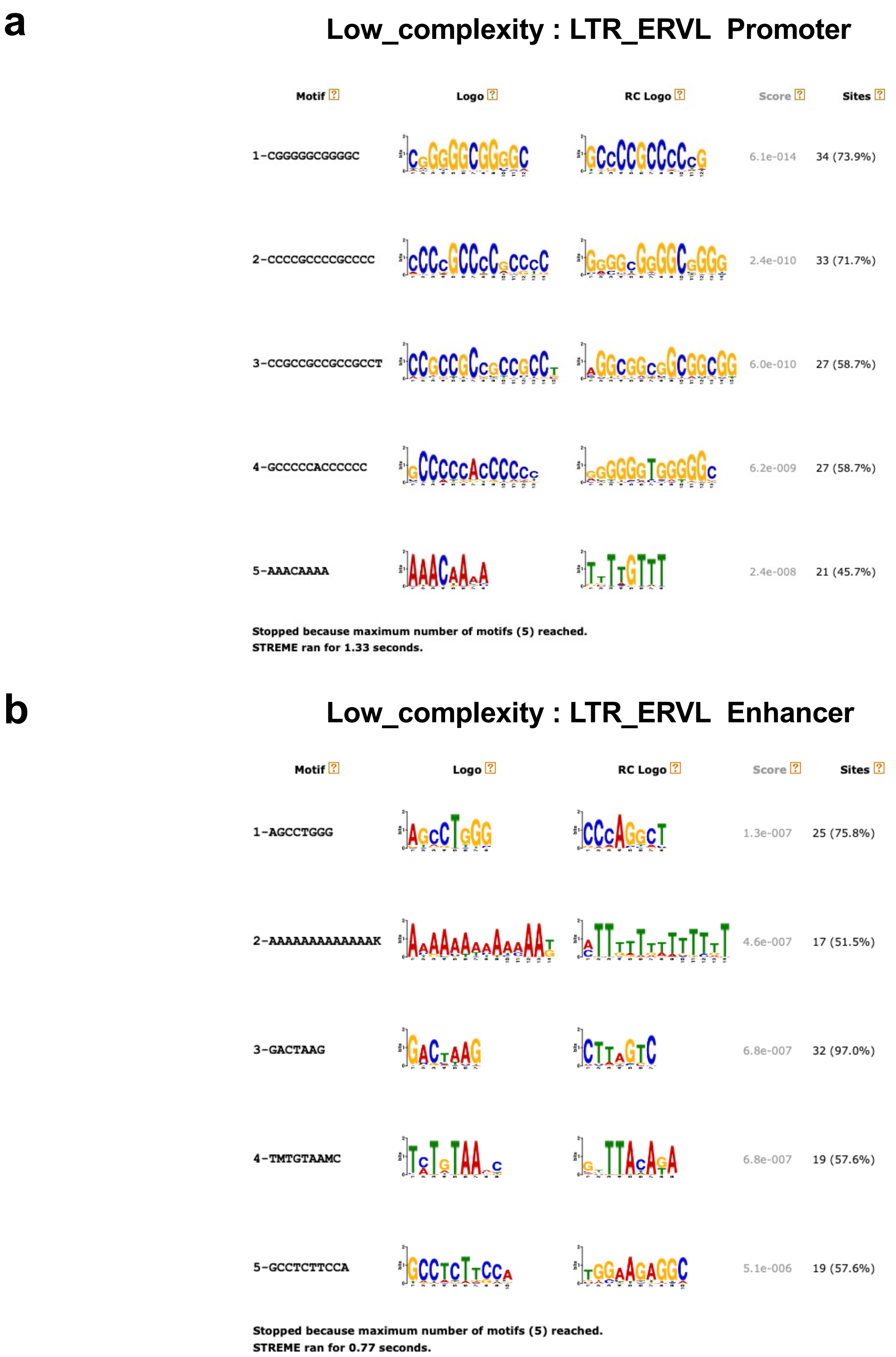


**Supplementary Figure S10: Motif discovery in Low_complexity : LTR_ERVL repeat–repeat interaction pairs in LTED cells. a**, *De novo* motif enrichment analysis of promoter-associated low_complexity regions from Low_complexity : LTR_ERVL REEPIs enriched in LTED cells. **b**, Motif enrichment analysis of enhancer-associated LTR_ERVL regions from the same interactions.

#### Supplementary Figure S10

Organism: *Homo sapiens*

SP1 *[ENSP00000329357]*

Transcription factor Sp1; Transcription factor that can activate or repress transcription in response to physiological and pathological stimuli. Binds with high affinity to GC-rich motifs and regulates the expression of a large number of genes involved in a variety of processes such as cell growth, apoptosis, differentiation and immune responses. Highly regulated by post-translational modifications (phosphorylations, sumoylation, proteolytic cleavage, glycosylation and acetylation). Binds also the PDGFR-alpha G-box promoter. May have a role in modulating the cellular response to DNA da [...]

*See SP1 functional neighbourhood network*

↔

SMAD3 *[ENSP00000332973]*

Mothers against decapentaplegic homolog 3; Receptor-regulated SMAD (R-SMAD) that is an intracellular signal transducer and transcriptional modulator activated by TGF-beta (transforming growth factor) and activin type 1 receptor kinases. Binds the TRE element in the promoter region of many genes that are regulated by TGF-beta and, on formation of the SMAD3/SMAD4 complex, activates transcription. Also can form a SMAD3/SMAD4/JUN/FOS complex at the AP- 1/SMAD site to regulate TGF-beta-mediated transcription. Has an inhibitory effect on wound healing probably by modulating both growth and m [...]

*See SMAD3 functional neighbourhood network*

Combined confidence of the functional interaction: 0.978 (very high)

EVIDENCE TABLE

| | | |
|---|---|---|
| Neighborhood in the Genome: | none / insignificant. | |
| Gene Fusions: | none / insignificant | |
| Cooccurrence Across Genomes: | none / insignificant | |
| Co-Expression: | yes (score 0.056). | show |
| Experimental/Biochemical Data: | yes (score 0.820). In addition, putative homologs were found interacting in other organisms (score 0.071). | show |
| Association in Curated Databases: | yes (score 0.500). | show |
| Co-Mentioned in Pubmed Abstracts: | yes (score 0.772). In addition, putative homologs are mentioned together in other organisms (score 0.051). | show |

**Supplementary Figure S11: STRING-based interaction evidence between SP1 and SMAD3.** Protein–protein interaction network analysis from the STRING database indicates a high-confidence interaction between SP1 and SMAD3 (combined score = 0.978).

### Supplementary Figure S11

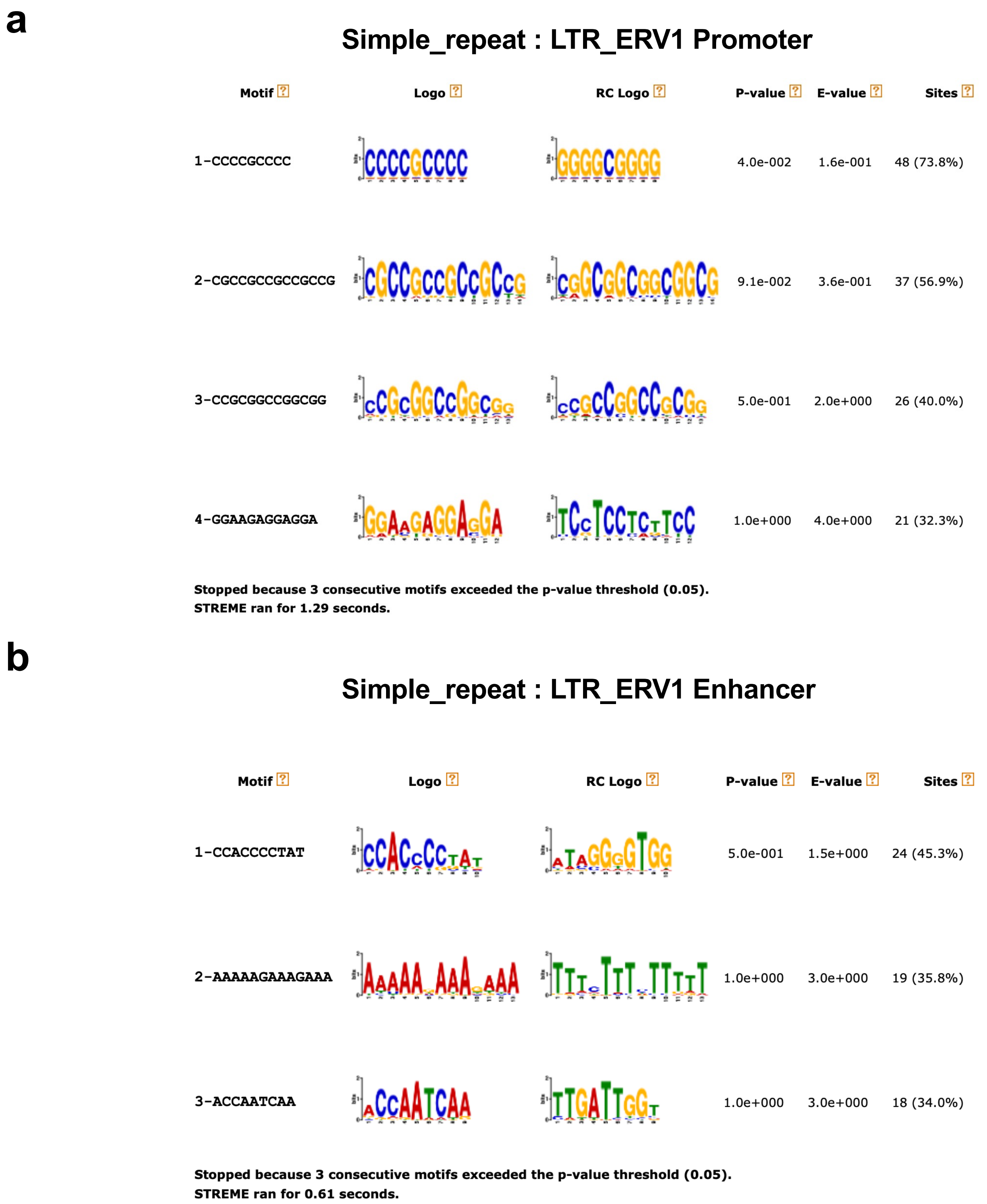


**Supplementary Figure S12: Motif discovery in Simple_repeat : LTR_ERV1 repeat–repeat interaction pairs in LTED cells. a**, *De novo* motif enrichment analysis of promoter-associated Simple_repeat regions within LTED-enriched Simple_repeat : LTR_ERV1 REEPIs. **b**, Motif enrichment analysis of enhancer-associated LTR_ERV1 regions from the same REEPIs.

#### Supplementary Figure S12

Organism: *Homo sapiens*

NFYC *[ENSP00000312617]*

Nuclear transcription factor Y subunit gamma; Component of the sequence-specific heterotrimeric transcription factor (NF-Y) which specifically recognizes a 5'-CCAAT-3' box motif found in the promoters of its target genes. NF-Y can function as both an activator and a repressor, depending on its interacting cofactors.

*See NFYC functional neighbourhood network*

↔

SP1 *[ENSP00000329357]*

Transcription factor Sp1; Transcription factor that can activate or repress transcription in response to physiological and pathological stimuli. Binds with high affinity to GC-rich motifs and regulates the expression of a large number of genes involved in a variety of processes such as cell growth, apoptosis, differentiation and immune responses. Highly regulated by post-translational modifications (phosphorylations, sumoylation, proteolytic cleavage, glycosylation and acetylation). Binds also the PDGFR-alpha G-box promoter. May have a role in modulating the cellular response to DNA da [...]

*See SP1 functional neighbourhood network*

Combined confidence of the functional interaction: 0.704 (high)

EVIDENCE TABLE

| | | |
|---|---|---|
| Neighborhood in the Genome: | none / insignificant. | |
| Gene Fusions: | none / insignificant | |
| Cooccurrence Across Genomes: | none / insignificant | |
| Co-Expression: | yes (score 0.155). | show |
| Experimental/Biochemical Data: | yes (score 0.120). In addition, putative homologs were found interacting in other organisms (score 0.095). | show |
| Association in Curated Databases: | yes (score 0.500). | show |
| Co-Mentioned in Pubmed Abstracts: | yes (score 0.246). In addition, putative homologs are mentioned together in other organisms (score 0.054). | show |

**Supplementary Figure S13: STRING-based interaction evidence between SP1 and NFYC.** Protein–protein interaction analysis using the STRING database shows a high-confidence interaction between SP1 and NFYC (combined score = 0.704).

### Supplementary Figure S13

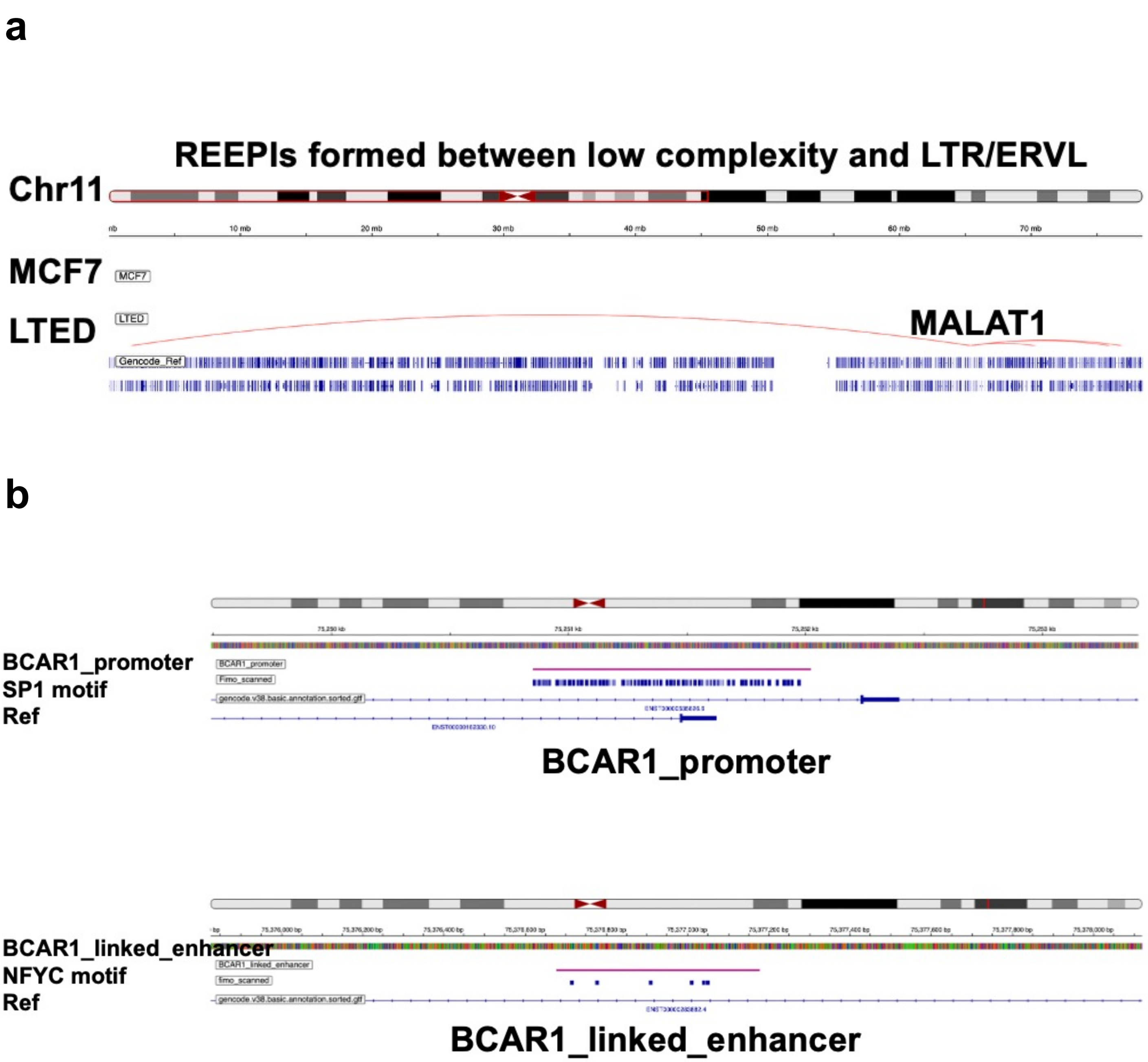


**Supplementary Figure S14: Representative REEPIs showing transcription factor–associated repeat pairings at endocrine resistance–related gene loci. a,** Genome browser visualization of a representative Low_complexity : LTR/ERVL REEPIs in LTED cells overlapping the *MALAT1* promoter region. The interaction arc (red) indicates a repeat-mediated enhancer–promoter contact that is absent in MCF7 cells. **b,** Motif inspection of the Simple_repeat : LTR/ERV1 REEPIs connecting the *BCAR1* promoter and its linked enhancer in LTED cells. The promoter anchor contains SP1-binding motifs, while the paired enhancer anchor harbors NFYC-binding motifs.

#### Supplementary Figure S14

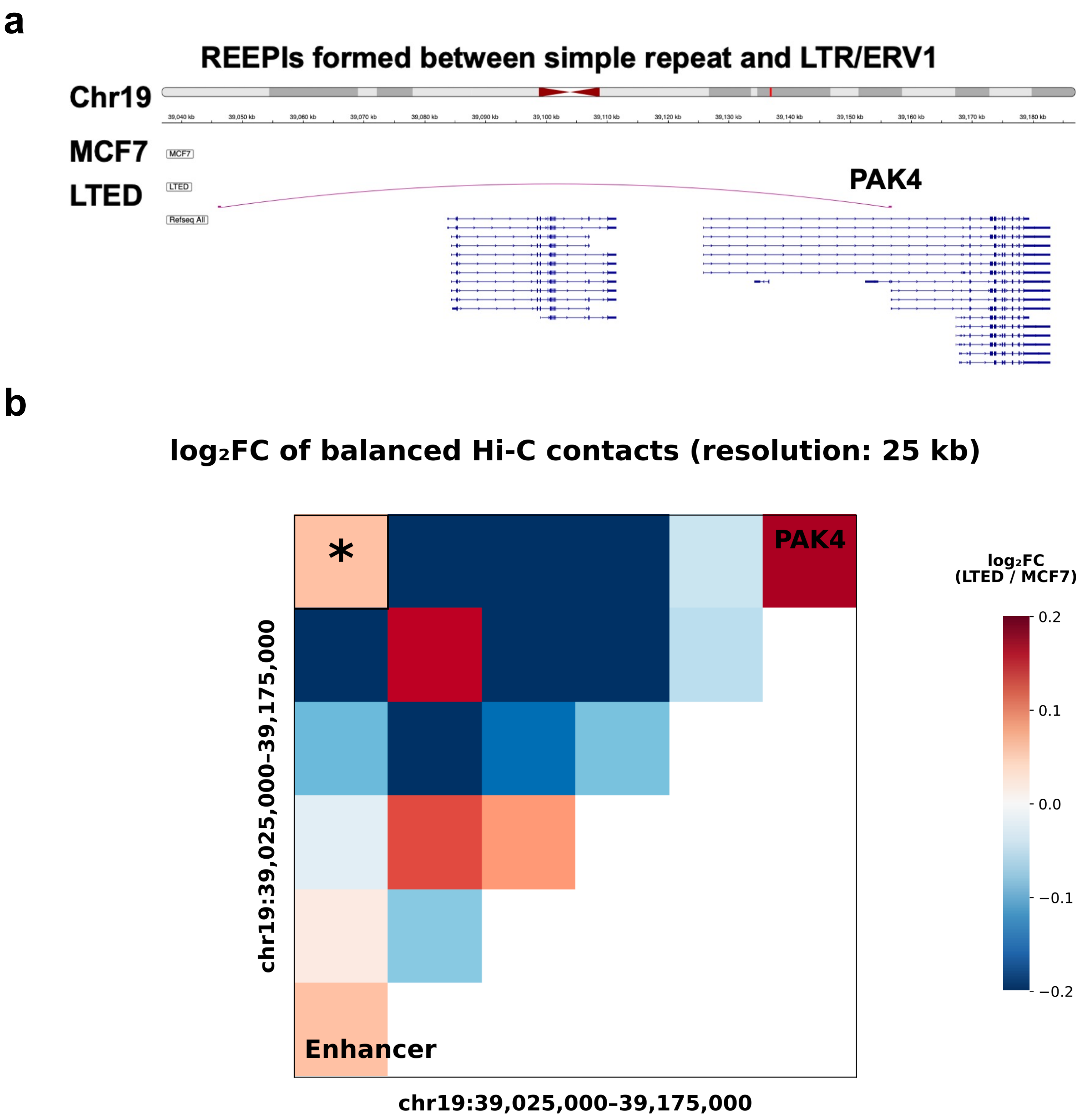


**Supplementary Figure S15: *PAK4*-associated LTED-specific REEPIs and local Hi-C contact change. a,** genome-browser view of a Simple_repeat : LTR/ERV1 REEPI connecting the *PAK4* promoter with a linked enhancer in LTED cells. The interaction is detected in LTED but not in MCF7 cells. **b,** heat map showing $\log_2$ fold changes in balanced Hi-C contacts between LTED and MCF7 cells at the *PAK4* locus (LTED/MCF7; 25-kb resolution; chr19:39,025,000–39,175,000). The asterisk marks the bin pair containing the *PAK4* promoter and the linked enhancer. Positive values indicate higher balanced contact signals in LTED cells.

#### Supplementary Figure S15
