## Supplementary File 1 for "Repeat element–anchored enhancer–promoter interactions (REEPIs) shape the regulatory landscape of hormone-resistant breast cancer"

### STREME

Sensitive, Thorough, Rapid, Enriched Motif Elicitation

For further information on how to interpret these results please access https://meme-suite.org/meme/doc/streme.html.
To get a copy of the MEME software please access https://meme-suite.org.

If you use STREME in your research, please cite the following paper:
Timothy L. Bailey, "STREME: accurate and versatile sequence motif discovery", *Bioinformatics*, Mar. 24, 2021. [full text]

Discovered Motifs | Inputs & Settings | Program Information | Motifs in MEME Text Format | Matching Sequences | Matching Sites | Results in XML Format

#### Description

LTED_P

#### Discovered Motifs

Next Top

| Motif | Logo | RC Logo | P-value | E-value | Sites | More | Submit/ Download | Positional Distribution | Matches per Sequence |
|---|---|---|---|---|---|---|---|---|---|
| 1-CCCCGCCCCCRCC | 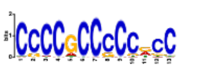 | 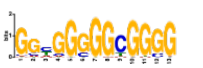 | 7.5e-021 | 7.0e-019 | 1561 (41.3%) | | | 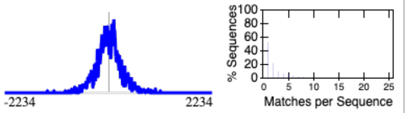 | |
| 2-GCHGCCGCCGCCGCC | 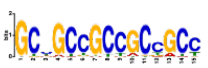 | 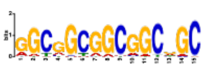 | 2.2e-015 | 2.1e-013 | 808 (21.4%) | | | 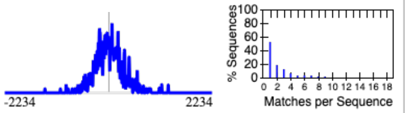 | |
| 3-CCCSGCCCCGCCCC | 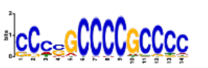 | 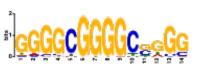 | 7.4e-015 | 6.8e-013 | 1262 (33.4%) | | | 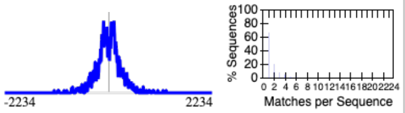 | |
| 4-AAAAAAAAAAAAAAA | 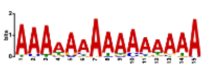 | 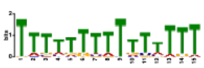 | 3.8e-010 | 3.6e-008 | 563 (14.9%) | | | 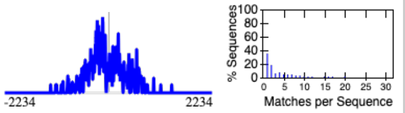 | |
| 5-CCTCCTCCTCCTCC | 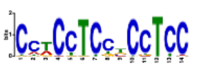 | 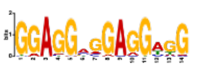 | 1.3e-009 | 1.2e-007 | 653 (17.3%) | | | 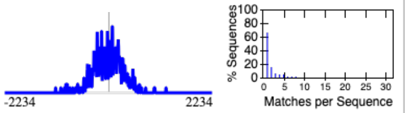 | |
| 6-CGCATGCGCAGW | 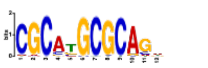 | 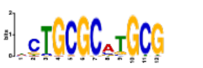 | 2.2e-008 | 2.0e-006 | 360 (9.5%) | | | 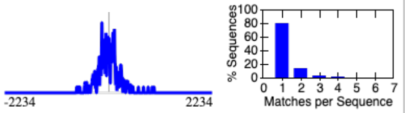 | |
| 7-CAGCCAATCA | 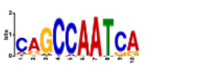 | 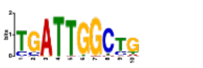 | 6.9e-008 | 6.4e-006 | 426 (11.3%) | | | 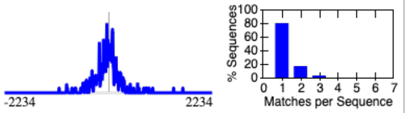 | |
| 8-AGGGAAGAGGMAG | 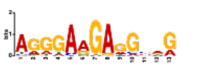 | 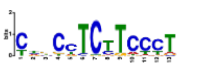 | 7.3e-007 | 6.8e-005 | 845 (22.3%) | | | 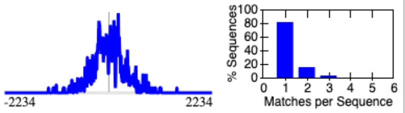 | |
| 9-GTGACGTCAC | 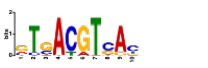 | 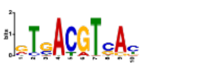 | 1.0e-006 | 9.4e-005 | 508 (13.4%) | | | 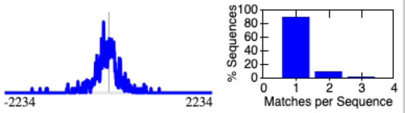 | |
| 10-TTTAAAATAHAA | 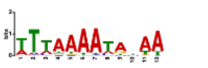 | 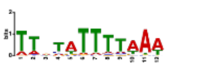 | 1.2e-006 | 1.1e-004 | 664 (17.5%) | | | 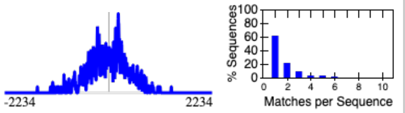 | |
| 11-AGGCTGGAGTGCAGT | 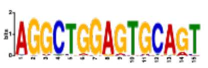 | 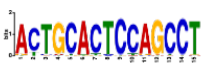 | 2.9e-006 | 2.7e-004 | 178 (4.7%) | | | 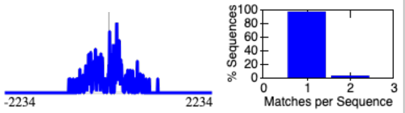 | |

**Stopped because 3 consecutive motifs exceeded the p-value threshold (0.05).**
**STREME ran for 4384.38 seconds.**

| Motif | Logo / RC Logo | P-value | E-value | Sites | More | Submit/ Download | Positional Distribution / Matches per Sequence |
|---|---|---|---|---|---|---|---|
| 12-RACTACAAYTCCCA | 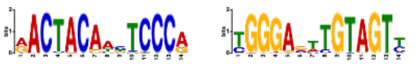 | 3.8e-006 | 3.5e-004 | 212 (5.6%) | | | 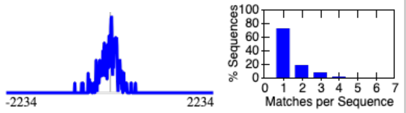 |
| 13-CRGCCCCGCCC | 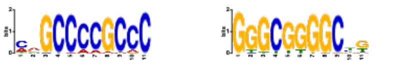 | 5.9e-006 | 5.5e-004 | 786 (20.8%) | | | 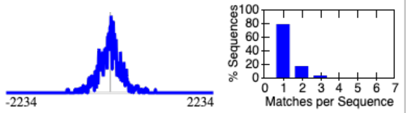 |
| 14-CCTCCCTCCCTCCC | 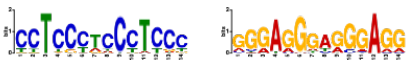 | 7.0e-006 | 6.5e-004 | 587 (15.5%) | | | 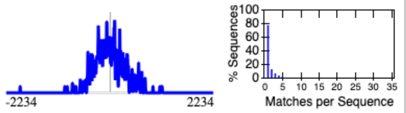 |
| 15-GAGAATCGCTTGAAC | 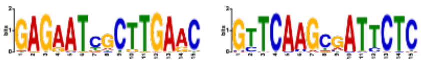 | 7.6e-006 | 7.1e-004 | 140 (3.7%) | | | 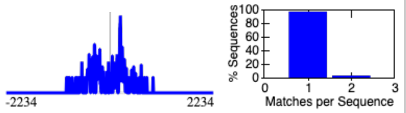 |
| 16-GCCTGTAATCCCAG | 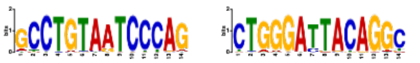 | 7.6e-006 | 7.1e-004 | 198 (5.2%) | | | 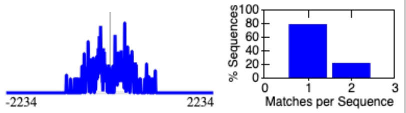 |
| 17-ATCTCGGCTCACTG | 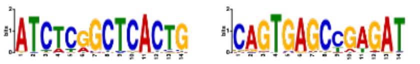 | 7.6e-006 | 7.1e-004 | 150 (4.0%) | | | 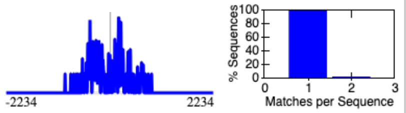 |
| 18-CAASATGGCG | 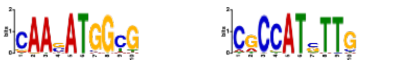 | 7.8e-006 | 7.3e-004 | 550 (14.5%) | | | 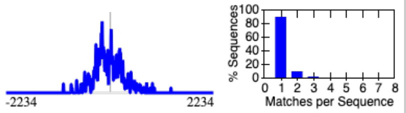 |
| 19-CAGCAGCAGCA | 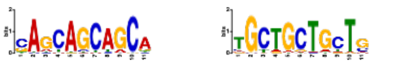 | 9.6e-006 | 8.9e-004 | 544 (14.4%) | | | 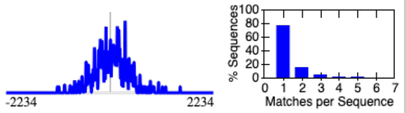 |
| 20-GCCTCAGYYTCCC | 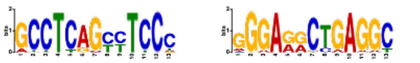 | 3.3e-005 | 3.1e-003 | 327 (8.6%) | | | 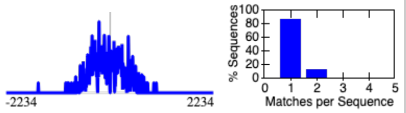 |
| 21-TATTTAYATA | 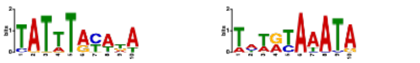 | 5.6e-005 | 5.2e-003 | 348 (9.2%) | | | 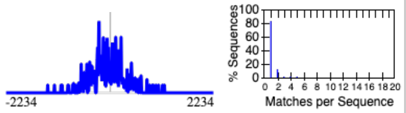 |
| 22-CCCCGCCTCCC | 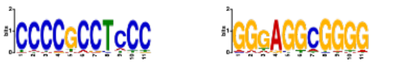 | 8.6e-005 | 8.0e-003 | 906 (23.9%) | | | 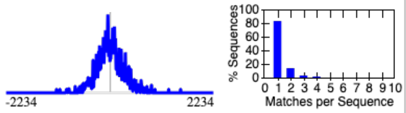 |
| 23-GCGCATGCGC | 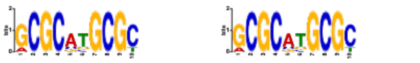 | 9.1e-005 | 8.5e-003 | 292 (7.7%) | | | 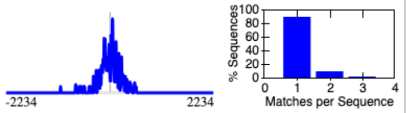 |
| 24-CCCGCCCCCGCCC | 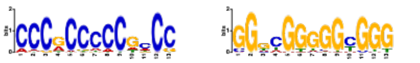 | 1.1e-004 | 9.9e-003 | 737 (19.5%) | | | 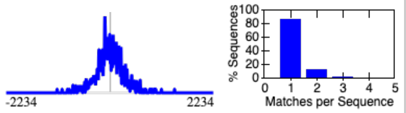 |
| 25-CCCCGCCCCTCBCC | 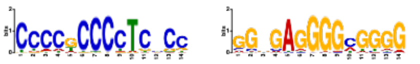 | 1.1e-004 | 1.0e-002 | 860 (22.7%) | | | 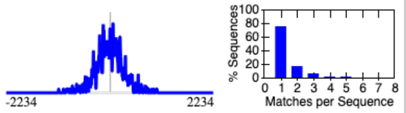 |

**Stopped because 3 consecutive motifs exceeded the p-value threshold (0.05).**
**STREME ran for 4384.38 seconds.**

| Motif | Logo | RC Logo | P-value | E-value | Sites | More | Submit/ Download | Positional Distribution | Matches per Sequence |
|---|---|---|---|---|---|---|---|---|---|
| 26-AGTTCGAGACCAGCC | | | 1.2e-004 | 1.1e-002 | 129 (3.4%) | | | | |
| 27-AAATAACAACAT | | | 2.4e-004 | 2.3e-002 | 730 (19.3%) | | | | |
| 28-CACACACACAC | | | 3.6e-004 | 3.4e-002 | 243 (6.4%) | | | | |
| 29-CGCCCCCTCCCC | | | 4.3e-004 | 4.0e-002 | 461 (12.2%) | | | | |
| 30-GGCGCGGTGGCTCAC | | | 4.9e-004 | 4.5e-002 | 114 (3.0%) | | | | |
| 31-CCCCGCGCCCCGCGC | | | 5.1e-004 | 4.7e-002 | 401 (10.6%) | | | | |
| 32-CGCCCGCCCGCCC | | | 7.0e-004 | 6.5e-002 | 364 (9.6%) | | | | |
| 33-AGGTGAGTGC | | | 7.2e-004 | 6.7e-002 | 315 (8.3%) | | | | |
| 34-CAASTTG | | | 7.6e-004 | 7.1e-002 | 648 (17.1%) | | | | |
| 35-CCCCACCCCCA | | | 9.6e-004 | 8.9e-002 | 744 (19.7%) | | | | |
| 36-CACGTGACT | | | 1.1e-003 | 1.0e-001 | 698 (18.4%) | | | | |
| 37-CCCCACCC | | | 1.2e-003 | 1.1e-001 | 816 (21.6%) | | | | |
| 38-CGCCGCCGCCGSC | | | 1.3e-003 | 1.2e-001 | 820 (21.7%) | | | | |
| 39-CCCGCCCCCTC | | | 1.8e-003 | 1.7e-001 | 515 (13.6%) | | | | |

**Stopped because 3 consecutive motifs exceeded the p-value threshold (0.05).**
**STREME ran for 4384.38 seconds.**

| Motif | Logo / RC Logo | P-value | E-value | Sites | More | Submit/ Download | Positional Distribution / Matches per Sequence |
|---|---|---|---|---|---|---|---|
| 40-CATTTTACAKA | 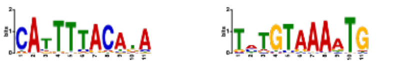 | 1.9e-003 | 1.7e-001 | 239 (6.3%) | | | 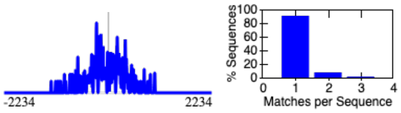 |
| 41-CCCAGCCCCG | 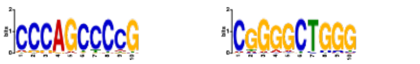 | 1.9e-003 | 1.7e-001 | 851 (22.5%) | | | 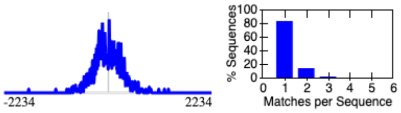 |
| 42-GGCTCCGCCCC | 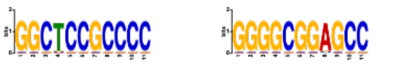 | 2.0e-003 | 1.8e-001 | 98 (2.6%) | | | 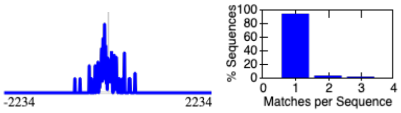 |
| 43-CCCGGGTCCCCA | 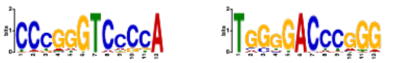 | 2.2e-003 | 2.0e-001 | 115 (3.0%) | | | 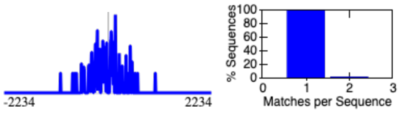 |
| 44-CTCCTCCTCCC | 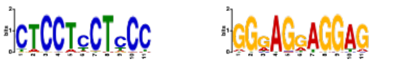 | 2.7e-003 | 2.5e-001 | 463 (12.2%) | | | 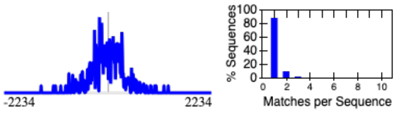 |
| 45-CCCCTCCCC | 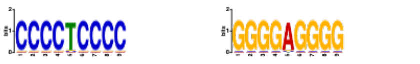 | 2.7e-003 | 2.5e-001 | 398 (10.5%) | | | 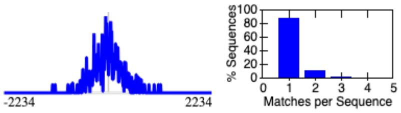 |
| 46-AAGATGAA | 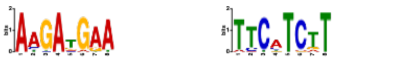 | 3.7e-003 | 3.4e-001 | 609 (16.1%) | | | 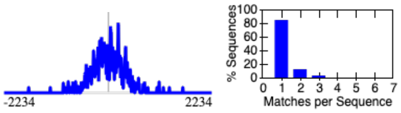 |
| 47-CCCGGCCCGGCCCGS | 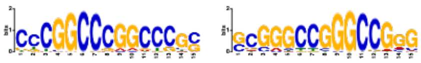 | 3.9e-003 | 3.6e-001 | 355 (9.4%) | | | 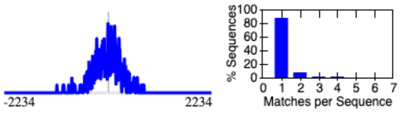 |
| 48-AAACCCCGTCTCT | 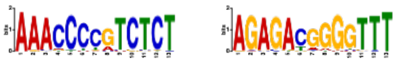 | 3.9e-003 | 3.6e-001 | 94 (2.5%) | | | 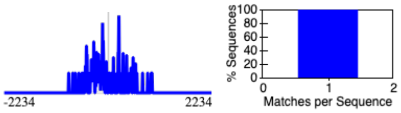 |
| 49-CGGAGCCCGAGCC | 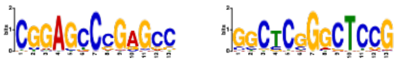 | 4.7e-003 | 4.4e-001 | 355 (9.4%) | | | 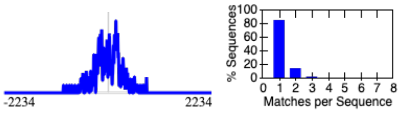 |
| 50-MGGAAGTGR | 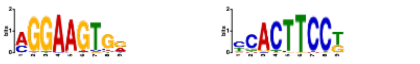 | 6.2e-003 | 5.8e-001 | 988 (26.1%) | | | 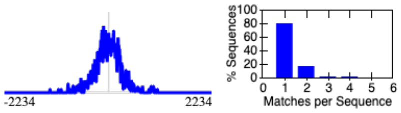 |
| 51-CGCCCGCCGCCCGCC | 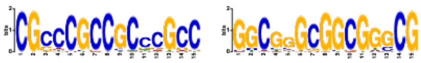 | 6.5e-003 | 6.0e-001 | 134 (3.5%) | | | 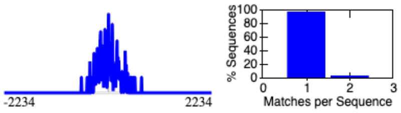 |
| 52-AGGTAAG | 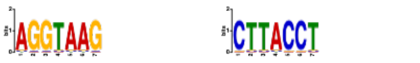 | 6.8e-003 | 6.4e-001 | 268 (7.1%) | | | 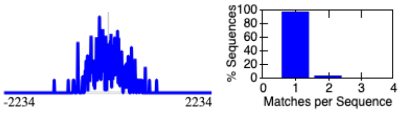 |
| 53-CCACCACCACCAC | 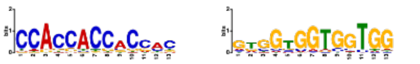 | 7.0e-003 | 6.5e-001 | 525 (13.9%) | | | 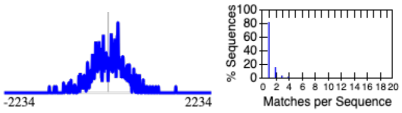 |

**Stopped because 3 consecutive motifs exceeded the p-value threshold (0.05).**
**STREME ran for 4384.38 seconds.**

| Motif | Logo | RC Logo | P-value | E-value | Sites | More | Submit/ Download | Positional Distribution | Matches per Sequence |
|---|---|---|---|---|---|---|---|---|---|
| 54-CTSATTGGCCC | 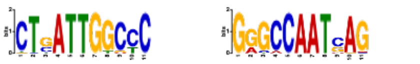 | | 7.8e-003 | 7.3e-001 | 59 (1.6%) | | | 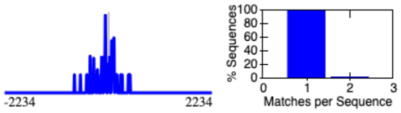 | |
| 55-CCCCGCCCC | 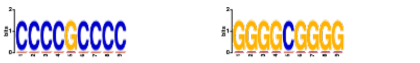 | | 8.2e-003 | 7.6e-001 | 338 (8.9%) | | | 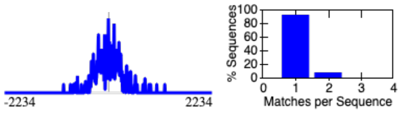 | |
| 56-GCGACAGAGCGAGAC | 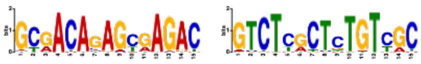 | | 1.1e-002 | 1.0e+000 | 141 (3.7%) | | | 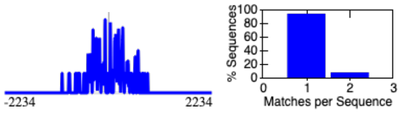 | |
| 57-CAGGTGAG | 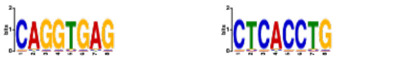 | | 1.2e-002 | 1.1e+000 | 261 (6.9%) | | | 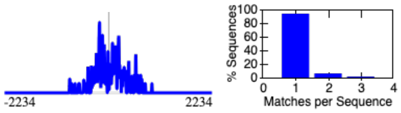 | |
| 58-CCCGGCTCC | 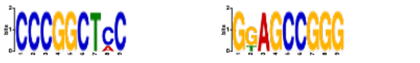 | | 1.2e-002 | 1.1e+000 | 408 (10.8%) | | | 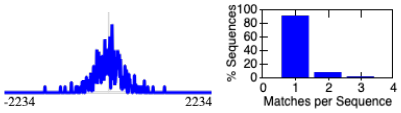 | |
| 59-GTGACTCA | 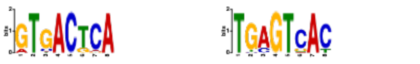 | | 1.2e-002 | 1.1e+000 | 402 (10.6%) | | | 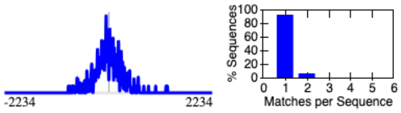 | |
| 60-ASCTGCTKCB | 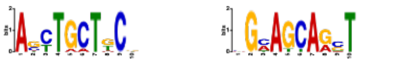 | | 1.5e-002 | 1.4e+000 | 369 (9.8%) | | | 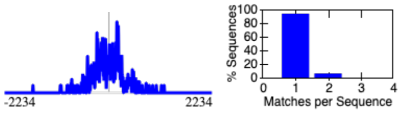 | |
| 61-CAASTTG | 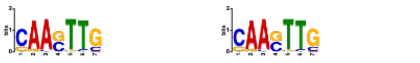 | | 1.6e-002 | 1.5e+000 | 775 (20.5%) | | | 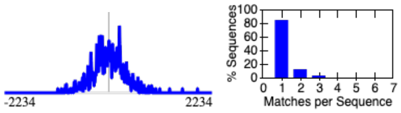 | |
| 62-CTCGCCCACCTC | 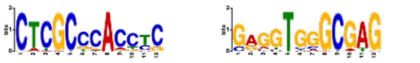 | | 1.8e-002 | 1.6e+000 | 125 (3.3%) | | | 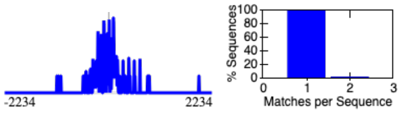 | |
| 63-CGCCTCCTCCT | 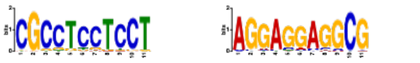 | | 1.8e-002 | 1.7e+000 | 223 (5.9%) | | | 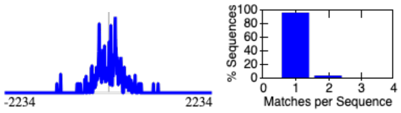 | |
| 64-MMCCACCCWCCC | 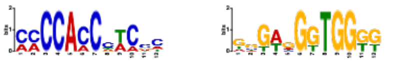 | | 1.8e-002 | 1.7e+000 | 419 (11.1%) | | | 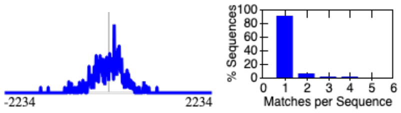 | |
| 65-GAAAGGAAGAA | 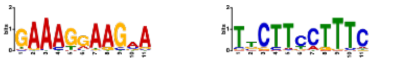 | | 2.2e-002 | 2.0e+000 | 379 (10.0%) | | | 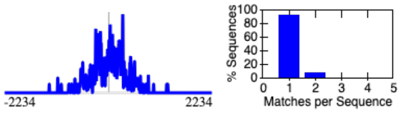 | |
| 66-CCGAGGCCG | 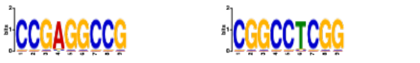 | | 2.2e-002 | 2.1e+000 | 288 (7.6%) | | | 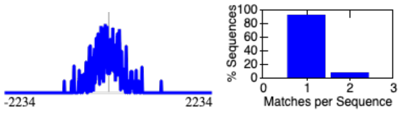 | |
| 67-AGAGAGAGAGAGAG | 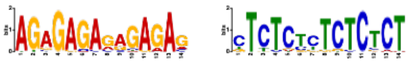 | | 2.5e-002 | 2.3e+000 | 185 (4.9%) | | | 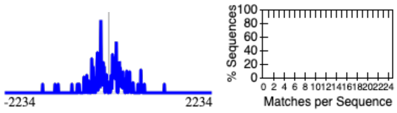 | |

**Stopped because 3 consecutive motifs exceeded the p-value threshold (0.05).**
**STREME ran for 4384.38 seconds.**

| Motif | Logo | RC Logo | P-value | E-value | Sites | More | Submit/ Download | Positional Distribution | Matches per Sequence |
|---|---|---|---|---|---|---|---|---|---|
| 68-CCCGGGTCTCGG | | | 2.8e-002 | 2.6e+000 | 400 (10.6%) | | | | |
| 69-GCCAATGGSAGG | | | 2.9e-002 | 2.7e+000 | 151 (4.0%) | | | | |
| 70-CCAAGGTCAC | | | 2.9e-002 | 2.7e+000 | 256 (6.8%) | | | | |
| 71-CGCGGGCCGGGGGCG | | | 3.1e-002 | 2.9e+000 | 70 (1.8%) | | | | |
| 72-CCCCTCCCCCCGCC | | | 3.1e-002 | 2.9e+000 | 69 (1.8%) | | | | |
| 73-ATCACCTGAGGTC | | | 3.1e-002 | 2.9e+000 | 67 (1.8%) | | | | |
| 74-ACACACAC | | | 3.2e-002 | 3.0e+000 | 173 (4.6%) | | | | |
| 75-AAGTTTG | | | 3.3e-002 | 3.0e+000 | 225 (5.9%) | | | | |
| 76-CTCGCTCGCTCSCT | | | 3.3e-002 | 3.0e+000 | 146 (3.9%) | | | | |
| 77-CCGGGGCTGCDG | | | 3.5e-002 | 3.2e+000 | 712 (18.8%) | | | | |
| 78-GCGGCCCCTTTAA | | | 3.5e-002 | 3.3e+000 | 85 (2.2%) | | | | |
| 79-CCATGGCRACC | | | 3.5e-002 | 3.3e+000 | 115 (3.0%) | | | | |
| 80-GCSGCSGCSGC | | | 3.6e-002 | 3.3e+000 | 709 (18.7%) | | | | |
| 81-AAGGAAAKAAAA | | | 3.8e-002 | 3.5e+000 | 173 (4.6%) | | | | |

**Stopped because 3 consecutive motifs exceeded the p-value threshold (0.05).**
**STREME ran for 4384.38 seconds.**

| Motif | Logo | RC Logo | P-value | E-value | Sites | More | Submit/ Download | Positional Distribution | Matches per Sequence |
|---|---|---|---|---|---|---|---|---|---|
| 82-CCCGCGCCGCC | | | 3.8e-002 | 3.6e+000 | 121 (3.2%) | | | | |
| 83-ACCAATCAVA | | | 3.8e-002 | 3.6e+000 | 152 (4.0%) | | | | |
| 84-GCTGCAGC | | | 4.2e-002 | 3.9e+000 | 1123 (29.7%) | | | | |
| 85-AATGAATGA | | | 4.3e-002 | 4.0e+000 | 333 (8.8%) | | | | |
| 86-AAACAAAG | | | 4.3e-002 | 4.0e+000 | 643 (17.0%) | | | | |
| 87-ATGCAAATA | | | 4.4e-002 | 4.1e+000 | 281 (7.4%) | | | | |
| 88-ACCAATGGGGA | | | 4.7e-002 | 4.3e+000 | 169 (4.5%) | | | | |
| 89-CCCGGGACCCCG | | | 4.9e-002 | 4.6e+000 | 214 (5.7%) | | | | |
| 90-ACCCCGAC | | | 4.9e-002 | 4.6e+000 | 231 (6.1%) | | | | |
| 91-CCTCTGCCTCT | | | 1.1e-001 | 1.1e+001 | 202 (5.3%) | | | | |
| 92-AGGTGAAGATG | | | 1.2e-001 | 1.1e+001 | 440 (11.6%) | | | | |
| 93-AACTTCAACCAA | | | 1.5e-001 | 1.4e+001 | 239 (6.3%) | | | | |

**Stopped because 3 consecutive motifs exceeded the p-value threshold (0.05).**
**STREME ran for 4384.38 seconds.**

#### Inputs & Settings

Previous Next Top

##### Sequences

| Role | Source | Alphabet | Sequence Count | Total Size |
|---|---|---|---|---|
| Positive (primary) Sequences | LTED_RADICL_links_Cis_with_distance_removed_CNV_promoter.bed.fa | DNA | 3784 | 3223014 |
| Negative (control) Sequences | 2-Order Shuffled Positive Sequences | DNA | 3784 | 3199362 |

##### Background Model

**Source:** built from the negative (control) sequences

**Order:** 2 (only order-0 shown)

| Name | Freq. | Bg. | Bg. | Freq. | Name |
|---|---|---|---|---|---|

| Name | Freq. | Bg. | | | | Bg. | Freq. | Name |
|---|---|---|---|---|---|---|---|---|
| Adenine | 0.174 | 0.174 | A | ~ | T | 0.174 | 0.174 | Thymine |
| Cytosine | 0.326 | 0.326 | C | ~ | G | 0.326 | 0.326 | Guanine |

##### Other Settings

| | |
|---|---|
| **Strand Handling** | Both the given and reverse complement strands are processed. |
| **Objective Function** | Differential Enrichment |
| **Statistical Test** | Binomial Test |
| **Minimum Motif Width** | 6 |
| **Maximum Motif Width** | 15 |
| **Sequence Shuffling** | Negative sequences are positives shuffled preserving 3-mer frequencies. |
| **Test Set** | 10% of the input sequences were randomly assigned to the test set. |
| **Word Evaluation** | Up to 25 words of each width from 6 to 15 were evaluated to find seeds. |
| **Seed Refinement** | Up to 4 seeds of each width from 6 to 15 were further refined. |
| **Refinement Iterations** | Up to 20 iterations were allowed when refining a seed. |
| **Random Number Seed** | 0 |
| **Trimming of Control Sequences** | Trimming of control sequences was allowed. |
| **Total Length** | The total length of each sequence set was limited to 4.00e+6. |
| **Maximum Motif p-value** | Stop when the p-value is greater than 0.05 for 3 consecutive motifs. |
| **Maximum Motifs to Find** | No maximum number of motifs. |
| **Maximum Run Time** | 14400 seconds. |

Previous Top

**STREME version**
5.5.8 (Release date: Thu May 15 15:01:46 2025 -0700)

**Reference**
Timothy L. Bailey, "STREME: accurate and versatile sequence motif discovery", *Bioinformatics*, Mar. 24, 2021. [full text]

**Command line**
```
streme --verbosity 1 --oc . --dna --totallength 4000000 --time 14400 --minw 6 --maxw 15 --thresh 0.05 --align center --dfile description --p
LTED_RADICL_links_Cis_with_distance_removed_CNV_promoter.bed.fa
```
