## Supplementary File 2 for "Repeat element–anchored enhancer–promoter interactions (REEPIs) shape the regulatory landscape of hormone-resistant breast cancer"

### STREME

Sensitive, Thorough, Rapid, Enriched Motif Elicitation

For further information on how to interpret these results please access https://meme-suite.org/meme/doc/streme.html.
To get a copy of the MEME software please access https://meme-suite.org.

If you use STREME in your research, please cite the following paper:
Timothy L. Bailey, "STREME: accurate and versatile sequence motif discovery", *Bioinformatics*, Mar. 24, 2021. [full text] If you use STREME in your research, please cite the following paper:
Timothy L. Bailey, "STREME: accurate and versatile sequence motif discovery", *Bioinformatics*, Mar. 24, 2021. [full text]

DISCOVERED MOTIFS | INPUTS & SETTINGS | PROGRAM INFORMATION | MOTIFS IN MEME TEXT FORMAT | MATCHING SEQUENCES | MATCHING SITES | RESULTS IN XML FORMAT

#### DESCRIPTION

LTED_e

#### DISCOVERED MOTIFS

Next Top

| Motif | Logo | RC Logo | P-value | E-value | Sites | More | Submit/ Download | Positional Distribution | Matches per Sequence |
|---|---|---|---|---|---|---|---|---|---|
| 1-AAAAAAAAAAAAA | 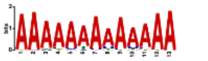 | 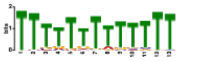 | 2.0e-008 | 8.6e-007 | 411 (16.7%) | | | 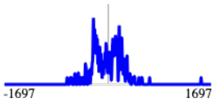 | 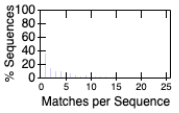 |
| 2-CAGCCTGGGCRACA | 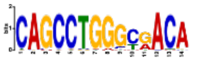 | 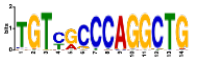 | 2.1e-006 | 9.2e-005 | 198 (8.1%) | | | 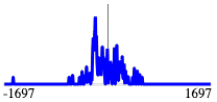 | 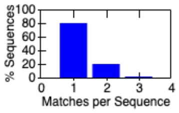 |
| 3-GCCTGTAATCCCAGC | 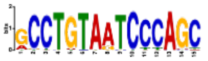 | 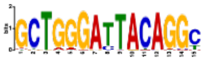 | 8.3e-006 | 3.6e-004 | 180 (7.3%) | | | 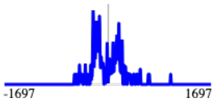 | 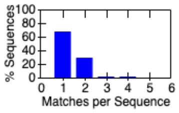 |
| 4-RTGASTCAY | 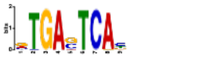 | 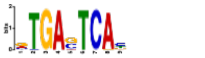 | 2.3e-005 | 1.0e-003 | 326 (13.3%) | | | 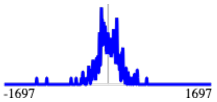 | 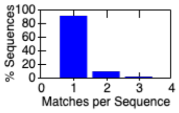 |
| 5-GCCACCRCGCCCGGC | 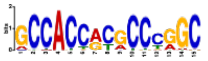 | 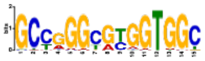 | 3.3e-005 | 1.4e-003 | 149 (6.1%) | | | 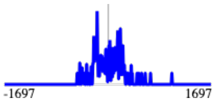 | 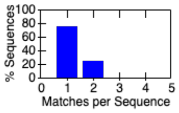 |
| 6-GCCTCAGCCTCC | 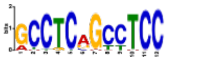 | 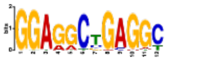 | 5.7e-005 | 2.5e-003 | 290 (11.8%) | | | 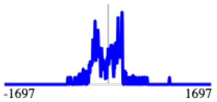 | 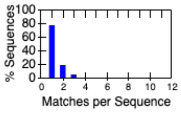 |
| 7-CCCCACCCTCC | 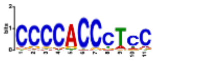 | 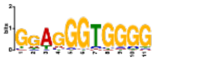 | 1.1e-004 | 4.8e-003 | 720 (29.3%) | | | 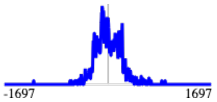 | 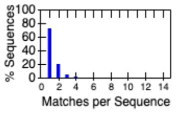 |
| 8-GGAGAATCGCTTGA | 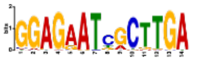 | 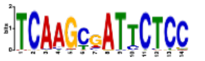 | 1.3e-004 | 5.7e-003 | 142 (5.8%) | | | 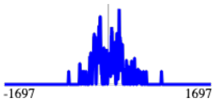 | 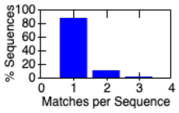 |
| 9-GCCCCGCCCC | 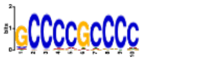 | 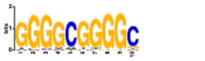 | 1.4e-004 | 6.3e-003 | 464 (18.9%) | | | 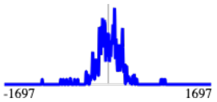 | 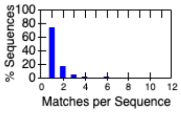 |
| 10-GAGTGCAGTGGCGCG | 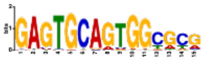 | 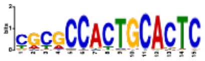 | 1.5e-004 | 6.5e-003 | 145 (5.9%) | | | 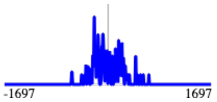 | 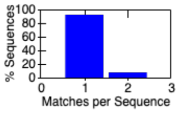 |

**Stopped because 3 consecutive motifs exceeded the p-value threshold (0.05).**
**STREME ran for 963.16 seconds.**

| Motif | Logo | RC Logo | P-value | E-value | Sites | More | Submit/ Download | Positional Distribution | Matches per Sequence |
|---|---|---|---|---|---|---|---|---|---|
| 11-GAGACGGRGTYTCRC | | | 2.2e-004 | 9.6e-003 | 181 (7.4%) | | | | |
| 12-CCCCACCCCCACC | | | 2.3e-004 | 1.0e-002 | 556 (22.6%) | | | | |
| 13-TCTCGGCTCACTGCA | | | 2.8e-004 | 1.2e-002 | 136 (5.5%) | | | | |
| 14-AAATATTTATT | | | 1.8e-003 | 8.0e-002 | 211 (8.6%) | | | | |
| 15-ATGTTGGCCAGGMTG | | | 2.0e-003 | 9.0e-002 | 72 (2.9%) | | | | |
| 16-GTAAAYAR | | | 2.4e-003 | 1.1e-001 | 447 (18.2%) | | | | |
| 17-ATGCAAACA | | | 2.9e-003 | 1.3e-001 | 398 (16.2%) | | | | |
| 18-AGGGCTGGGG | | | 3.8e-003 | 1.7e-001 | 400 (16.3%) | | | | |
| 19-GGCRGATCACCTGA | | | 4.1e-003 | 1.8e-001 | 43 (1.8%) | | | | |
| 20-TATTTTTAAA | | | 4.4e-003 | 1.9e-001 | 190 (7.7%) | | | | |
| 21-CTCGAACTCCTGACC | | | 6.8e-003 | 3.0e-001 | 134 (5.5%) | | | | |
| 22-AATAAATA | | | 7.8e-003 | 3.4e-001 | 477 (19.4%) | | | | |
| 23-CTCCTYCTCCCTCC | | | 1.1e-002 | 4.7e-001 | 295 (12.0%) | | | | |
| 24-AAAGAAAGAAAA | | | 1.2e-002 | 5.1e-001 | 383 (15.6%) | | | | |

**Stopped because 3 consecutive motifs exceeded the p-value threshold (0.05).**
**STREME ran for 963.16 seconds.**

| Motif | Logo | RC Logo | P-value | E-value | Sites | More | Submit/ Download | Positional Distribution | Matches per Sequence |
|---|---|---|---|---|---|---|---|---|---|
| 25-ACCCGGGAGGCRGAG | | | 1.6e-002 | 7.1e-001 | 70 (2.9%) | | | | |
| 26-CCGGCCCTGCCC | | | 2.0e-002 | 8.8e-001 | 109 (4.4%) | | | | |
| 27-AGGCGGGGCTGG | | | 2.3e-002 | 1.0e+000 | 282 (11.5%) | | | | |
| 28-CAAGTC | | | 2.5e-002 | 1.1e+000 | 569 (23.2%) | | | | |
| 29-GTGTGTGTGTGTR | | | 2.8e-002 | 1.2e+000 | 287 (11.7%) | | | | |
| 30-CCATGGCAACG | | | 3.2e-002 | 1.4e+000 | 52 (2.1%) | | | | |
| 31-CAGCAGCAGCAGC | | | 3.2e-002 | 1.4e+000 | 64 (2.6%) | | | | |
| 32-GAGAGAGACAGAG | | | 3.2e-002 | 1.4e+000 | 66 (2.7%) | | | | |
| 33-ACTTTGGGAGGCCRA | | | 3.2e-002 | 1.4e+000 | 60 (2.4%) | | | | |
| 34-CCTCCCCGCC | | | 3.3e-002 | 1.5e+000 | 159 (6.5%) | | | | |
| 35-TAATAATAATAM | | | 3.6e-002 | 1.6e+000 | 57 (2.3%) | | | | |
| 36-GGGGAGGGGAGGGGA | | | 3.8e-002 | 1.7e+000 | 240 (9.8%) | | | | |
| 37-CTTCCTTCT | | | 4.0e-002 | 1.8e+000 | 606 (24.7%) | | | | |
| 38-ACTTCCTG | | | 4.1e-002 | 1.8e+000 | 144 (5.9%) | | | | |

**Stopped because 3 consecutive motifs exceeded the p-value threshold (0.05).**
**STREME ran for 963.16 seconds.**

| Motif | Logo | RC Logo | P-value | E-value | Sites | More | Submit/ Download | Positional Distribution | Matches per Sequence |
|---|---|---|---|---|---|---|---|---|---|
| 39-GCCGCCGCCGCC | 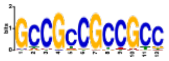 | 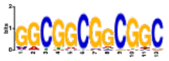 | 4.1e-002 | 1.8e+000 | 181 (7.4%) | | | 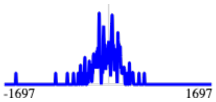 | 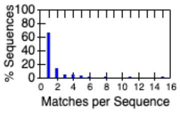 |
| 40-CTCCTCCCCATC | 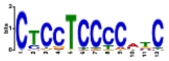 | 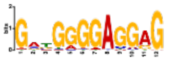 | 4.4e-002 | 1.9e+000 | 453 (18.5%) | | | 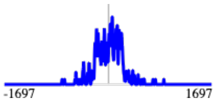 | 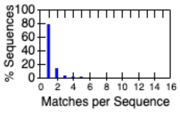 |
| 41-GATGACGA | 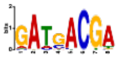 | 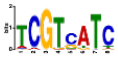 | 4.6e-002 | 2.0e+000 | 197 (8.0%) | | | 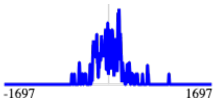 | 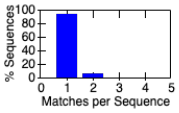 |
| 42-CCCCGCCCGCCCSC | 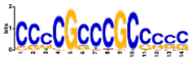 | 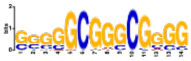 | 1.6e-001 | 6.8e+000 | 117 (4.8%) | | | 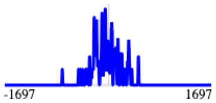 | 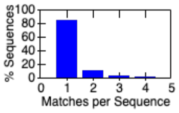 |
| 43-CAGCCCYGGGSHC | 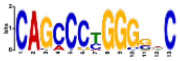 | 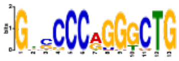 | 1.9e-001 | 8.4e+000 | 59 (2.4%) | | | 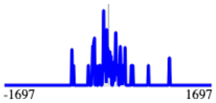 | 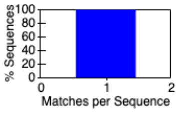 |
| 44-GGGAGGGAGGGA | 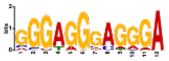 | 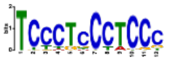 | 6.7e-001 | 2.9e+001 | 158 (6.4%) | | | 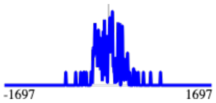 | 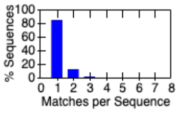 |

**Stopped because 3 consecutive motifs exceeded the p-value threshold (0.05).**
**STREME ran for 963.16 seconds.**

#### Inputs & Settings

Previous Next Top

##### Sequences

| Role | Source | Alphabet | Sequence Count | Total Size |
|---|---|---|---|---|
| Positive (primary) Sequences | LTED_RADICL_links_Cis_with_distance_removed_CNV_enhancer.bed.fa | DNA | 2455 | 1538389 |
| Negative (control) Sequences | 2-Order Shuffled Positive Sequences | DNA | 2455 | 1536916 |

##### Background Model

**Source:** built from the negative (control) sequences

**Source:** built from the negative (control) sequences

**Order:** 2 (only order-0 shown)

| Name | Freq. | Bg. | | | | Bg. | Freq. | Name |
|---|---|---|---|---|---|---|---|---|
| Adenine | 0.222 | 0.222 | A | ~ | T | 0.222 | 0.222 | Thymine |
| Cytosine | 0.278 | 0.278 | C | ~ | G | 0.278 | 0.278 | Guanine |

| Name | Freq. | Bg. | | | | Bg. | Freq. | Name |
|---|---|---|---|---|---|---|---|---|
| Adenine | 0.222 | 0.222 | A | ~ | T | 0.222 | 0.222 | Thymine |
| Cytosine | 0.278 | 0.278 | C | ~ | G | 0.278 | 0.278 | Guanine |

##### Other Settings

| | |
|---|---|
| **Strand Handling** | Both the given and reverse complement strands are processed. |
| **Objective Function** | Differential Enrichment |
| **Statistical Test** | Binomial Test |
| **Minimum Motif Width** | 6 |
| **Maximum Motif Width** | 15 |
| **Sequence Shuffling** | Negative sequences are positives shuffled preserving 3-mer frequencies. |
| **Test Set** | 10% of the input sequences were randomly assigned to the test set. |
| **Word Evaluation** | Up to 25 words of each width from 6 to 15 were evaluated to find seeds. |
| **Seed Refinement** | Up to 4 seeds of each width from 6 to 15 were further refined. |
| **Refinement Iterations** | Up to 20 iterations were allowed when refining a seed. |
| **Random Number Seed** | 0 |
| **Trimming of Control Sequences** | Trimming of control sequences was allowed. |
| **Total Length** | The total length of each sequence set was limited to 4.00e+6. |
| **Maximum Motif p-value** | Stop when the p-value is greater than 0.05 for 3 consecutive motifs. |
| **Maximum Motifs to Find** | No maximum number of motifs. |
| **Maximum Run Time** | 14400 seconds. |

Previous Top

**STREME version**
5.5.8 (Release date: Thu May 15 15:01:46 2025 -0700)

**Reference**
Timothy L. Bailey, "STREME: accurate and versatile sequence motif discovery", *Bioinformatics*, Mar. 24, 2021. [full text]
**Reference**
Timothy L. Bailey, "STREME: accurate and versatile sequence motif discovery", *Bioinformatics*, Mar. 24, 2021. [full text]

**Command line**

```
streme --verbosity 1 --oc . --dna --totallength 4000000 --time 14400 --minw 6 --maxw 15 --thresh 0.05 --align center --dfile description --p
LTED_RADICL_links_Cis_with_distance_removed_CNV_enhancer.bed.fa
```
